# Extracellular vesicles from alternatively activated macrophages remodel the injured hepatic niche and promote repair

**DOI:** 10.64898/2026.11.17.767294

**Authors:** Melisande L Addison, Alastair M Kilpatrick, Maria Elena Candela, Candice Ashmore-Harris, Jennifer Cartwright, Rhona E Aird, Tak-Yung Man, Karen Ching, Despoina Kesidou, Justyna Cholewa-Waclaw, Christopher Humphries, Laura Wagstaff, James W Dear, Stuart J Forbes

## Abstract

Alternatively activated macrophages (AAMs) are in clinical development for acute liver injury, but their mechanisms of action remain incompletely understood. AAM therapy remodels the injured hepatic niche, promoting reparative macrophage states, suppressing inflammatory myeloid programmes and restoring hepatocyte metabolic and biosynthetic programmes. We identify extracellular vesicles released by AAMs (AAM-EVs) as essential mediators of AAM therapeutic activity. AAM-EVs enhance macrophage phagocytosis and induce reparative transcriptional and proteomic responses, including increased TREM2 abundance. Following administration in established APAP injury, AAM-EVs reduce necrosis and neutrophilic inflammation, and promote hepatocellular regeneration, and we observe preferential and rapid association of AAM-EVs with hepatic macrophages alongside reparative phenotypic shift. Moreover, pharmacological inhibition of EV release attenuates parent AAM efficacy and TREM2⁺ macrophage accumulation, while EVs derived from Dicer-deficient AAMs exhibit reduced therapeutic activity, implicating miRNA-associated mechanisms. Comparative single-nucleus transcriptomics reveals overlapping but distinct macrophage and hepatocyte responses to AAMs and AAM-EVs. Together, these findings identify EV-mediated communication as an important mechanism of macrophage-induced liver repair and support the development of AAM-EVs as a cell-free therapeutic strategy.

**Highlights:**

- Alternatively activated macrophage (AAM) therapy remodels endogenous macrophage and hepatocyte states during liver repair
- AAM-derived EVs enhance phagocytosis and promote repair in APAP-ALI
- Inhibition of EV release attenuates AAM efficacy in APAP-ALI and prevents TREM2⁺ macrophage accumulation
- Dicer-deficient AAM-EVs show reduced therapeutic activity in acute liver injury
- AAMs and AAM-EVs share overlapping yet distinct transcriptomic profiles

## Introduction

Acetaminophen (APAP) overdose remains a leading cause of acute liver injury worldwide. Although N-acetylcysteine is effective when administered early, its benefit declines substantially with delayed presentation, and no therapies exist to prevent progression from APAP-induced acute liver injury (APAP-ALI) to acute liver failure(*1*). Once acute liver failure develops, treatment is limited to supportive care or, in selected cases, orthotopic liver transplantation, which is constrained by donor availability and associated with significant morbidity and mortality(*2, 3*).

Alternatively activated macrophages (AAMs) represent a promising off-the-shelf cell therapy for APAP-ALI. Administration of AAMs to mice with established injury (16 h post-APAP) reduces hepatic necrosis, dampens inflammation, and promotes regeneration, addressing a major unmet clinical need(*4*). Notably, allogeneic AAMs retain efficacy following cryopreservation, supporting their development as a scalable therapeutic, and are currently under evaluation in a first-in-human trial (MAIL, ISRCTN12637839)(*5, 6*). In parallel, autologous macrophage therapy has demonstrated safety and efficacy in liver cirrhosis (MATCH trial, ISRCTN10368050), including improved transplant free survival following a single infusion (*7–9*).

Despite these advances, how therapeutic macrophages influence endogenous immune and parenchymal cells within the injured liver remains incompletely understood. Previous work has suggested that AAM efficacy involves interactions with the host immune system, raising the possibility that secreted mediators contribute to their reparative activity (*4*). Using single-nucleus RNA sequencing (snRNA-seq), we provide evidence for reprogramming of both immune and parenchymal compartments by AAM therapy, promoting reparative macrophage states and restoring hepatocyte metabolic function. The widespread changes observed across host macrophage and hepatocyte populations raised the question of how transferred AAMs communicate with cells of the injured liver. Extracellular vesicles (EVs) are nanoscale lipid bilayer-bound particles enriched in proteins, lipids, and small RNAs, and can influence recipient cell function (*10*). Therefore, we investigated EVs as potential mediators of AAM activity. Here, we tested whether AAM-derived EVs (AAM-EVs) reproduce the effects of their parent cells, and whether EV release and EV microRNA cargo are required for AAM efficacy in the injured liver niche.

## RESULTS

### Biodynamic effects of human AAMs (hAAMs) in APAP-ALI revealed by single-nucleus RNA sequencing

Single-nucleus RNA sequencing was performed on FACS-sorted liver nuclei from APAP-injured mice treated with 5×10⁶ hAAMs or PBS at 16 h post-injury and culled at 36h, alongside uninjured controls (Fig. S1A). Supervised annotation identified all major hepatic lineages (Fig. S1B–C). Consistent with the reduced necrosis and increased regeneration observed with AAM therapy(*4*), we found that hAAM treatment altered cellular composition, with increased proportions of endothelial cells (OR = 1.82, FDR = 1.8 × 10⁻¹⁶) and Kupffer cells (OR = 1.30, FDR = 9.9 × 10⁻³), and a reduction in *Anxa2*+ hepatocytes, which we hereafter refer to as injury-response hepatocytes (IR-Heps) (OR = 0.66, FDR = 1.2 × 10⁻⁵) compared with PBS treated control (Fig. S1C) (*11–13*).

Reclustering of myeloid nuclei identified six populations, including *Trem2* macrophages (*Gpnmb⁺Cd36⁺Lgals3⁺*), monocytes (*F13a1⁺Ccr2⁺Cx3cr1⁺*), Kupffer cells (*Clec4f⁺Timd4⁺Cd5l⁺*), cycling Kupffer cells and dendritic cell subsets (Fig. 1A, Fig. S1D– E). Minimal alignment to the human genome suggested that the sequenced nuclei were predominantly host-derived rather than transferred hAAMs. Exploratory analysis of pooled nuclei showed increased proportions of cycling KCs (OR 1.71, P = 0.039, FDR = 0.148) and *Trem2* macrophages (OR = 1.38, P = 0.048, FDR = 0.148) with hAAM therapy. Within the *Trem2* macrophage population, hAAM treatment was associated with increased *Gpnmb*, *Myo1e* and *Hexb* and reduced *Dab2*, *Apoe* and *Ccr5* expression at 36h after APAP and 20h after administration of hAAM (P < 0.05; Fig. 1B; Fig. S2A). Over-representation analysis (ORA) identified enrichment of pathways related to hyaluronan catabolism, stress-granule and P-body assembly, as well as NF-κB signalling in upregulated genes (Fig. 1C, Fig. S2B). Given the wealth of evidence for the involvement of TREM2 macrophages in liver repair(*14, 15*), and that *Gpnmb* is a marker of this population(*16*), we decided to investigate this gene candidate further. Though we were unable to accurately quantify GPNMB expression, immunofluorescence demonstrated GPNMB/TREM2 co-expression within areas of pericentral necrosis, mainly in CD11b+ monocytes rather than CLEC4F⁺ Kupffer cells, and the percentage of TREM2⁺ cells was significantly increased following AAM treatment (Fig. 1D-E, Fig. S2C-D). Overall, these findings suggested that AAM treatment is capable of selective remodelling of macrophage programmes associated with lipid handling, cytoskeletal organisation and lysosomal function.

**FIGURE 1.**
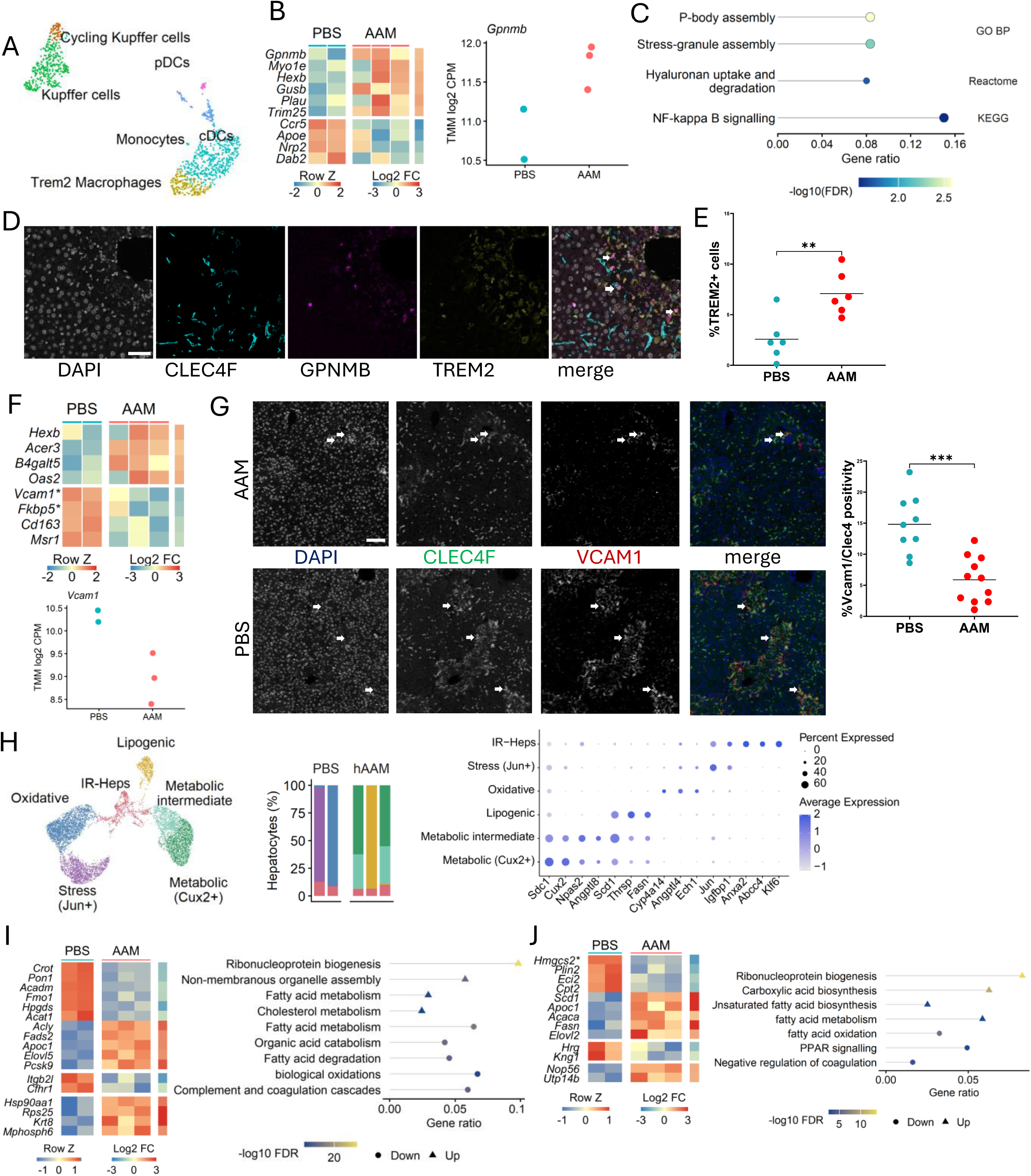
Treatment with hAAM therapy is associated with myeloid and hepatocyte reprogramming in the injured niche in APAP-ALI. (A) Reclustering of myeloid nuclei identifying distinct myeloid subsets (B) Heatmap showing key differentially expressed genes with scatter plot showing *Gpnmb* expression per animal in *Trem2* macrophages (C) Over-representation analysis of upregulated genes within *Trem2* macrophages using genes P < 0.05 and log2FC >0.5 with pathways relevant to APAP liver injury (D) Microscope images of immunofluorescent staining of liver sections for DAPI (grey), CLEC4F (cyan), GPNMB (magenta), and TREM2 (yellow), with white arrows highlighting GPNMB⁺TREM2⁺ co-localised cells in merged images (scale bar = 50µm). (E) Quantification scatter plot of TREM2/CD11b staining in APAP liver in mice given hAAM or PBS (representative images shown in supplementary) (F) Heatmap showing key differentially expressed genes with scatter plot (below) showing *Vcam1* expression per animal in KCs (G) Immunofluorescence staining of liver sections for DAPI (blue), CLEC4F (green) and VCAM1 (red), with white arrows highlighting CLEC4F⁺VCAM1⁺ co-localised cells in merged images, as well as a quantification scatter plot with % VCAM1/CLEC4F+ cells on the right (scale bar = 100 µm). (H) Hepatocyte reclustering showing UMAP with stacked bar chart showing proportion by animal, and dotplot of representative marker genes for each cluster (I) Representative heatmap and functional pathway analysis summary for differentially expressed genes in hepatocytes (excluding IR-Heps) by condition (J) Representative heatmap and functional pathway analysis summary for differentially expressed genes in IR-Heps by condition Heatmap colours indicate row-scaled normalised expression, with adjacent strips showing log₂ fold change for AAM versus PBS. Nominally significant gene candidates shown, with * indicating FDR <0.05. In the enrichment plots, horizontal position indicates gene ratio and colour indicates −log₁₀(pathway FDR); triangles and circles denote enrichment among increased and decreased genes, respectively.

The monocyte compartment showed a distinct treatment-associated transcriptional response, with reduced expression of genes associated with inflammatory monocyte recruitment and myeloid activation, including *Ccr2*, *Itgam*, *Cd74* and *Adam8*. Consistent with this, ORA identified reduced representation of programmes associated with myeloid-cell differentiation, phagocytosis and leukocyte proliferation. Conversely, hAAM treatment increased *Arg1* in monocytes together with an interferon-responsive programme, with enrichment of defence-response-to-virus and type II interferon-signalling pathways, which have previously been associated with uptake of extracellular vesicles from immune cell types(*17*) (Fig. S2A-B).

Kupffer cells similarly underwent transcriptional remodelling following hAAM treatment, with increased expression of *Hexb*, *Acer3*, *B4galt5* and *Oas2* and reduced expression of *Vcam1*, *Fkbp5*, *Cd1c3* and *Msr1* (Fig. 1F). Functional enrichment identified increased sphingolipid metabolism, driven by *Acer3*, *B4galt5*, *Hexb* and *Plpp3*, together with an interferon-associated antiviral programme. In contrast, pathways associated with scavenger-receptor-mediated ligand uptake were reduced, including decreased expression of *Cd1c3*, *Msr1* and *Stab2* (Fig. 1F; Fig. S2A–B). Immunofluorescence confirmed a reduction in VCAM1 positivity in CLEC4F⁺ Kupffer cells following hAAM treatment (Fig. 1G).

Reclustering of 7,734 hepatocyte nuclei from APAP-injured livers identified six transcriptional populations, including metabolic and lipogenic states following hAAM treatment, and stress-associated and oxidative states predominating in injured controls, with similar preponderance towards periportal zonation across conditions (Fig. 1H, S3A– C). We observed increased expression of lipid-metabolism gene candidates, including *Scd1*, *Fasn*, *Thrsp* and *Angptl8*. Enrichment analysis linked increased genes to fatty-acid biosynthesis and translation, and decreased genes to lipid catabolism and fatty-acid degradation (Fig. 1I), suggesting that hAAM treatment is associated with a shift towards hepatocyte biosynthetic programmes. We also observed hAAM-associated transcriptional changes in IR-Heps (Fig. 1J; Fig. S3D), characterised by increased expression of *Abcc4*, *Il34* and *Fosl1* and reduced expression of *Anxa3* and *Hmgcs2*. Within this population, enrichment analysis linked increased genes to biosynthetic processes and ribonucleoprotein-complex biogenesis, and decreased genes to fatty-acid oxidation and lipid catabolism (Fig. 1J), suggesting a shift towards normal hepatocyte function.

### AAM-derived extracellular vesicles promote reparative macrophage responses and phagocytosis

The widespread transcriptional remodelling of endogenous immune and parenchymal populations following hAAM therapy, coupled with the enrichment of known EV-uptake programmes, suggested that infused macrophages act beyond their immediate microenvironment through remote paracrine signalling. We therefore asked whether AAM-derived EVs contribute to the reparative effects of their parent cells. AAM-EVs were isolated from conditioned media by ultrafiltration and size exclusion chromatography. Parent AAMs showed typical morphology and expressed canonical alternative activation markers, including *Arg1*, *Chil3*, *Cd163* and *Mrc1* (Fig. S4B) (*18*). Nanoparticle tracking and transmission electron microscopy of isolated AAM-EVs demonstrated a mean particle diameter of ∼185 nm with typical EV morphology (Fig. S4C-D). Proteomic and immunoblot analyses detected canonical EV-associated proteins, including CD9, CD63, CD81, Flotillin and HSP70, with absence of GM130/GOLGA2 and GAPDH (Fig. S4D-E). AAM-EVs also retained expression of macrophage-associated proteins, including ARG1, CD206 and LGALS3 (Fig. S4E). PKH-26-labelled AAM-EVs were readily internalised by naïve BMDMs after a 3 h incubation, demonstrated by increased fluorescence relative to dye-only controls (Fig. S4F). Previous studies have reported the ability of macrophage EVs to modulate macrophage responses(*19, 20*). We observed *Arg1* to be a dose responsive marker of AAM-EV stimulation at baseline and under LPS conditions at doses 3 log folds lower than previously used in cell stimulation studies (10:1 and 100:1 EV-to-cell ratios, 5 ×10^6^ – 5×10^7^ particles/well). We also observed a dose response effect in secreted TNF and IL-10 (Fig. 2A–B).

**FIGURE 2.**
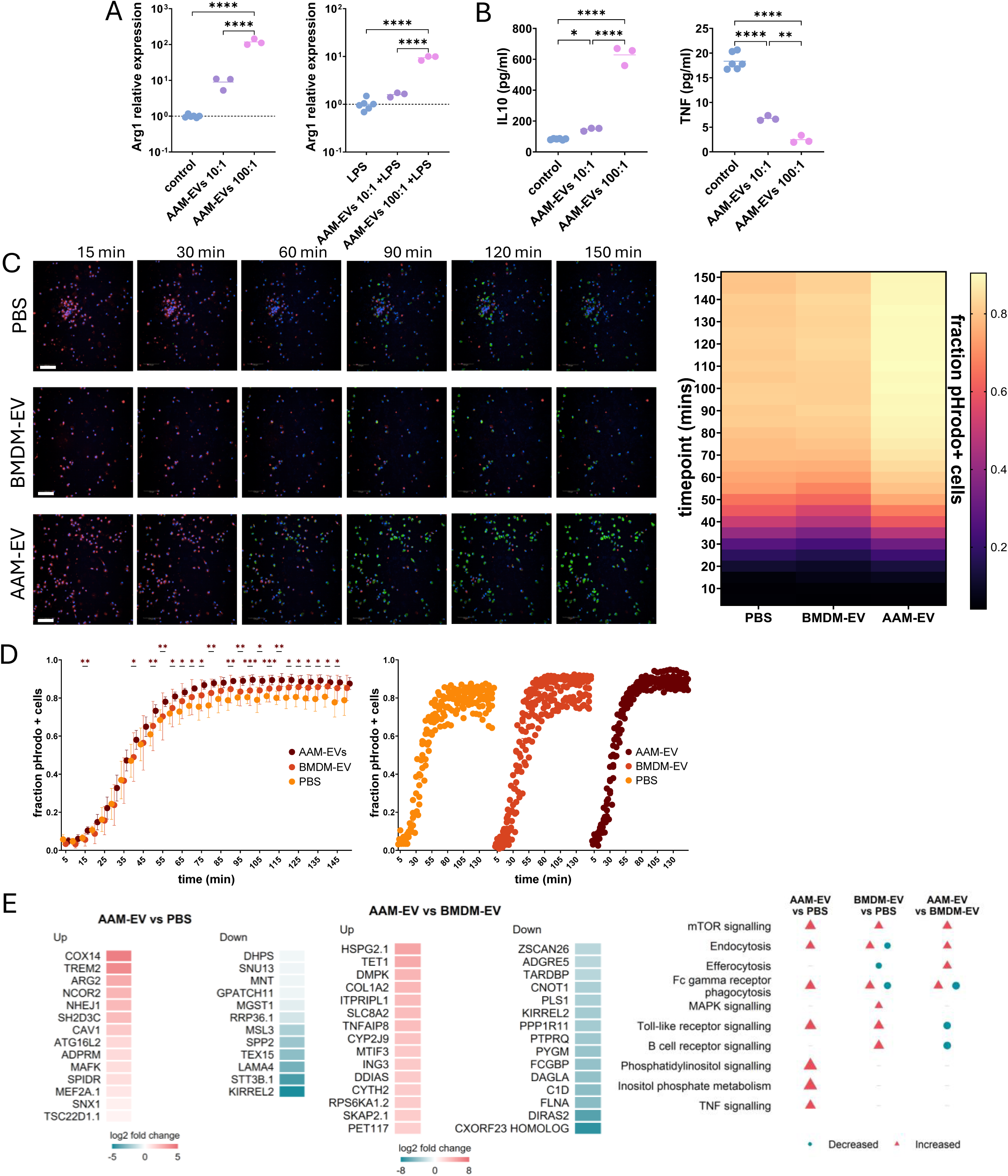
AAM-EVs promote a reparative phenotype in the context of inflammation and increase phagocytosis efficiency in naïve BMDMs, with proteomics revealing a rise in TREM2 expression. (A) qPCR analysis of *Arg1* gene expression in naïve BMDMs treated at AAM-EV-to-cell ratios of 10:1 or 100:1 for 16 h (left), as well as following LPS stimulation for 4h. Data are shown as fold change relative to unstimulated BMDMs (2^-ΔΔCT, log scale) (B) Measurement of cytokine levels in the media after naïve BMDMs were treated at AAM-EV-to-cell ratios of 10:1 or 100:1 for 16 h (pg/ml). (C) Representative images from real-time phagocytosis assays following addition of pHrodo-labelled particles to BMDMs treated with PBS, BMDM-EVs or AAM-EVs. CellMask Deep Red (red), NucBlue (blue) and intracellular pHrodo fluorescence (green) are shown. Heatmap illustrates the fraction of pHrodo-positive cells over time. Scale bars are 100μm. (D) Quantification of phagocytosis as the fraction of pHrodo-positive cells over time (mean +/−SD). The time course also plotted separately by treatment group for clarity (right), with each point representing an individual well (n = 6-8 wells per group). The fraction of pHrodo-positive cells was analysed using two-way repeated-measures ANOVA with treatment and time as factors, Geisser–Greenhouse correction for departures from sphericity and Dunnett’s multiple comparisons against PBS at each timepoint. Asterisks indicate significant differences between AAM-EVs and PBS; BMDM-EVs did not differ significantly from PBS. (E) Proteomic profiling of recipient BMDMs following treatment with AAM-EVs, BMDM-EVs or PBS. Heatmap shows log₂ fold changes for all significant differentially expressed proteins in BMDMs exposed to AAM-EVs vs those exposed to PBS for 24h (26 FDR-significant proteins, all displayed) and vs those exposed to BMDM-EVs (555 FDR-significant proteins, 15 shown in each direction). Exploratory KEGG over-representation analysis was performed separately on increased- and decreased-abundance protein lists defined by P < 0.05. Only pathways with BH-adjusted P < 0.05 are displayed. Coral triangles and teal circles indicate enrichment among increased and decreased proteins, respectively; symbol area represents fold enrichment, capped at 12.

To determine whether AAM-EVs could directly alter macrophage function, we performed a live-cell phagocytosis assay. AAM-EV–treated BMDMs phagocytosed pHrodo bioparticles more rapidly and to a greater extent than PBS-or BMDM-EV–treated controls (Fig. 2C–D). The fraction of pHrodo⁺ cells was significantly higher following AAM-EV treatment than PBS at the indicated timepoints, whereas BMDM-EVs did not differ significantly from PBS. We next performed dose–response studies using AAM-EVs ranging from 5 × 10^5^ to 5 × 10⁸ particles. Interestingly, AAM-EVs showed an inverse dose–response in this assay, with lower doses outperforming higher doses, possibly related to saturation of endocytosis/phagocytosis pathways at supraphysiological EV concentrations (Fig. S5A–B)(*17*).

Proteomic profiling of recipient BMDMs identified increased abundance of protein groups assigned to TREM2, ARG2 and COX14 following AAM-EV treatment compared with PBS (BH-adjusted FDR < 0.05; Fig. S5C–D). TREM2 is implicated in macrophage phagocytic function and tissue repair, while ARG2 and COX14 are involved in mitochondrial arginine metabolism and respiratory complex IV assembly, respectively, suggesting additional metabolic adaptation following EV treatment. KEGG over-representation analysis identified enrichment of endocytosis, Fcγ receptor-mediated phagocytosis, mTOR signalling and phosphatidylinositol signalling among increased-abundance proteins (Fig. 2E; Fig. S5C–D). Although both EV treatments elicited overlapping proteomic responses, direct comparison of AAM-EV- and BMDM-EV-treated macrophages identified enrichment of endocytosis, efferocytosis and mTOR signalling among increased-abundance proteins, and Toll-like receptor signalling among decreased-abundance proteins (pathway BH FDR < 0.05; Fig. 2E). Together, these findings indicate that AAM-EVs induce distinct changes in macrophage uptake-associated signalling and metabolic regulation, consistent with their greater phagocytic activity compared with BMDM-EVs.

### AAM-EVs reduce necrosis and inflammation and promote regeneration in APAP-ALI, exerting dose-dependent therapeutic effects

To determine whether AAM-EVs could reproduce the therapeutic effects of their parent cells in vivo, mice received PBS, 1 × 10⁶ AAMs or 1 × 10⁸ AAM-EVs at 16 h following APAP administration, when hepatic injury was already established. This delayed treatment window was selected to investigate therapeutic effects during established injury, when NAC is less effective (*21*) (Fig. 3A). The EV dose approximated the number of particles released by 1 × 10⁶ AAMs over 24 h, enabling comparison with the therapeutic parent-cell dose. Animals were assessed at 36 h after APAP administration. AAM-EV treatment significantly reduced serum ALT activity and hepatic necrosis compared with PBS-treated controls, together with reduced Ly6G⁺ neutrophil infiltration (Fig. 3B-C). We observed a trend towards increased Ki67+ and BRDU+ nuclei with AAM-EV treatment but observed no differences in TUNEL staining (Fig. 3C). Parent AAMs showed a similar directional reduction in hepatic necrosis and serum ALT and % change in ALT from 16h. This pilot study indicated that AAM-EVs are able to recapitulate key therapeutic effects of their parent cells and warranted further investigation.

**FIGURE 3.**
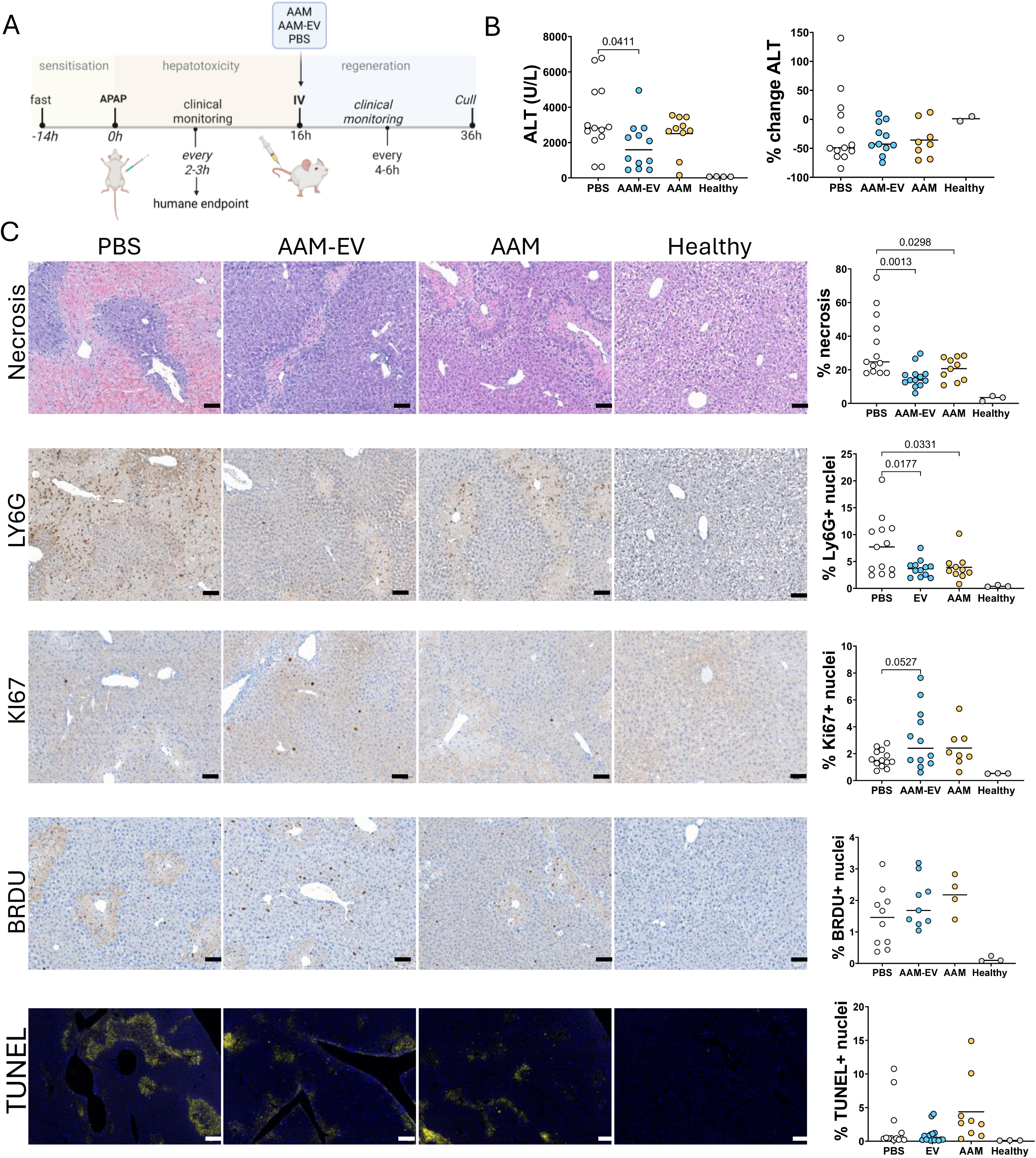
Intravenous AAM-EVs reduce necrosis and promote regeneration in mice with APAP-ALI. (A) Timeline of in vivo APAP experiment illustrating injection AAMs (1 × 10^6^, i.v.) and AAM-EVs (1 × 10^8^, i.v.) or PBS alone into APAP-ALI mice at 16 h, followed by clinical monitoring and cull at 36 h. (B) Serum ALT activity (U/L) and % change in ALT (16-36h) in APAP-ALI mice receiving PBS, AAMs, AAM-EVs or in healthy controls. (C) Representative images of liver HCE staining from APAP-ALI mice receiving indicated treatments with corresponding graph demonstrating quantification of percentage necrosis on the right. We also show Ly6G, Ki67, BrdU and TUNEL (respectively) positivity in mice receiving indicated treatments with quantification panel on the right. Scale bars 100µm. Ordinary ONE WAY ANOVA performed comparing PBS, AAM and AAM-EV groups. P values shown in graphs, n=8-13 per group.

We next investigated the therapeutic dose range of AAM-EVs, administering 1 × 10⁶–4 × 10⁸ particles (Fig. 4A). Based on EV yields from AAM cultures, 1 × 10⁸ particles was estimated to approximate the EV output over 24 h from the 1 × 10⁶ AAM dose used in previous therapeutic studies (*4*). As the dose-ranging experiments were performed across independent experimental cohorts with differences in baseline injury severity, individual-animal data were analysed using regression models incorporating experimental cohort as a categorical covariate. Hepatic necrosis, serum ALT and Ly6G staining were log-transformed, with log₁₀-transformed EV dose modelled as a continuous predictor; body-weight change was analysed on its original scale. Complementary categorical models were used to compare individual EV doses with PBS, with adjustment for multiple comparisons using Dunnett’s method (AAM treatment was not included in this analysis).

**FIGURE 4.**
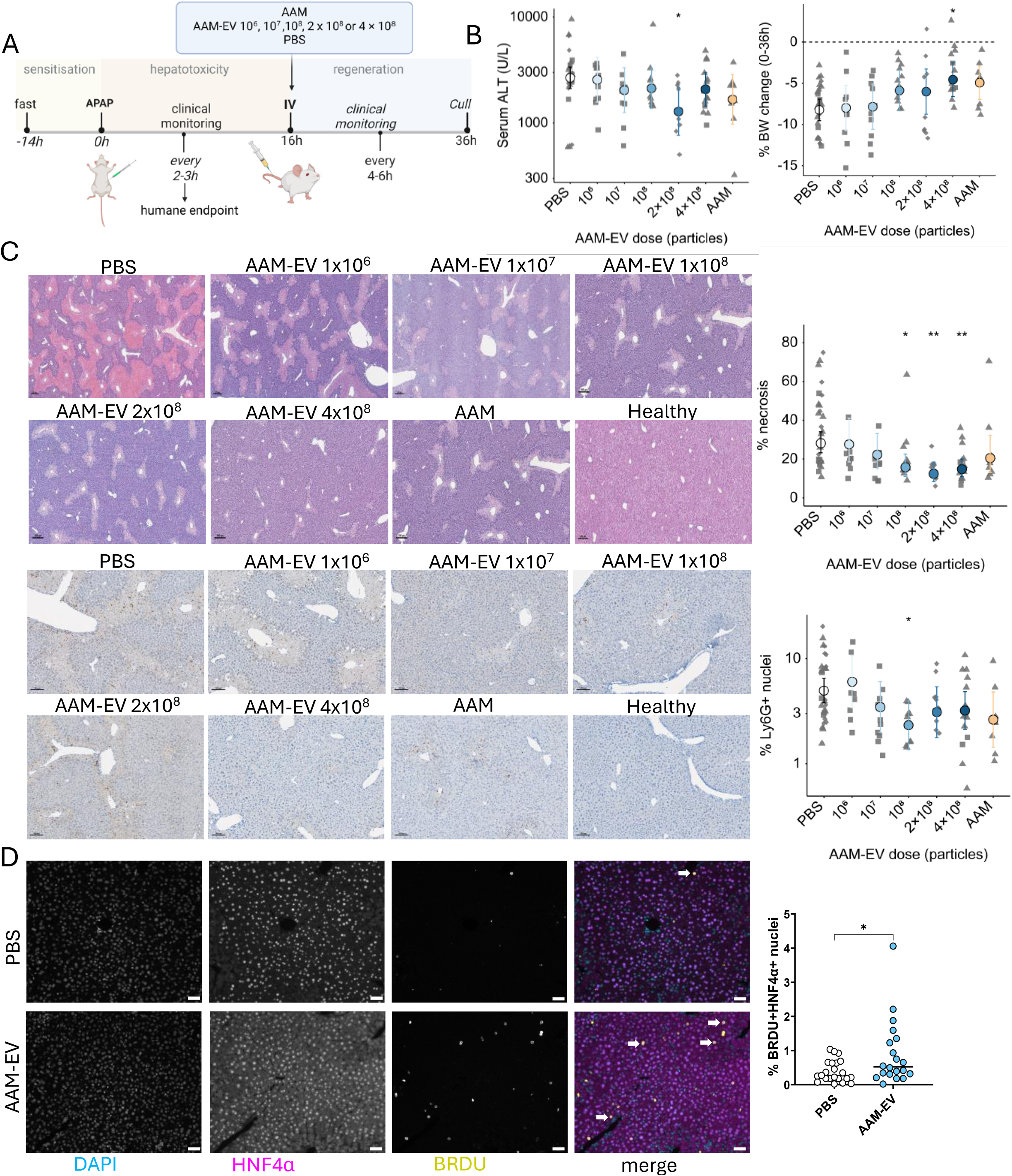
Dose dependent therapeutic effects of AAM-EVs in APAP-ALI. (A) Experimental timeline. Mice with established APAP-ALI were treated at 16 h with PBS, AAMs (1 × 10⁶ cells, i.v.) or the indicated dose of AAM-EVs and culled at 36 h. (B) Serum ALT activity (U/L) and percentage body weight change from 0–36 h following treatment with 1 × 10⁶, 1 × 10^7^, 1 × 10^8^, 2 x 10^8^ or 4 × 10⁸ AAM-EVs, 1 × 10⁶ AAMs or PBS. (C) Representative HCE (top panel) staining and Ly6G (bottom panel) immunopositivity with corresponding quantification panel following treatment with 1 × 10⁶ – 4 × 10⁸ AAM-EVs, AAMs or PBS (scale bars: HCE, 200 µm; Ly6G, 100 µm). (D) Dual immunofluorescence for BrdU (yellow) and HNF4α (magenta), with DAPI counterstain (cyan) in APAP-ALI mice treated with PBS or 4 × 10⁸ AAM-EVs. White arrows indicate BrdU⁺HNF4α⁺ hepatocytes; individual channels and merged images are shown, with BrdU quantification on the right. For dose-ranging analyses, individual animals are shown with symbols denoting experimental cohort (batch 1: diamonds, batch 2: triangles, batch 3: squares); coloured circles and error bars represent cohort-adjusted means and 95% CIs. Hepatic necrosis, ALT and Ly6G were log-transformed and body-weight change analysed on the original scale. Dose effects were assessed using cohort-adjusted regression, with individual doses compared with PBS using Dunnett-adjusted categorical models. AAMs are shown as a reference and were excluded from dose-response analyses. *P < 0.05, **P < 0.01, ***P < 0.001.

Increasing AAM-EV dose was associated with a significant reduction in hepatic necrosis (β = −0.244 per log₁₀ increase in dose, 95% CI −0.445 to −0.042, *P* = 0.019). In categorical analyses, doses of 1 × 10⁸, 2 × 10⁸ and 4 × 10⁸ particles significantly reduced hepatic necrosis relative to PBS controls (Dunnett-adjusted *P* = 0.016, 0.0012 and 0.0005, respectively), whereas lower doses did not. Secondary endpoints showed concordant treatment-associated changes, including a trend towards reduced body-weight loss with increasing AAM-EV dose (Fig. 4B–C).

Markers of regeneration further supported a reparative effect of AAM-EVs, with reduced P21 staining and increased BRDU incorporation (Fig. S6A–B). Immunofluorescent staining of HNF4α/ERG/BRDU demonstrated that this increase occurred predominantly within hepatocytes, indicating enhanced hepatocellular regeneration (Fig. 4D, Fig. S6C). Circulating cytokine levels were not significantly altered between treatment groups (Fig. S6D).

### AAM-EVs preferentially accumulate in injured liver and associate with hepatic macrophage populations

To determine whether AAM-EVs exhibit similar biodistribution to their parent cells, 4×10⁸ DiR-labelled AAM-EVs were injected into mice with established APAP-ALI at 16 h following APAP administration. In vivo imaging demonstrated significantly increased hepatic DiR signal at 6 h and 20 h post-injection compared with PBS-treated controls. Ex vivo organ imaging confirmed preferential accumulation within the liver, with additional signal detected in the spleen and lung at 20 h (Fig. S7A–B). To investigate intrahepatic EV uptake, biotin-labelled AAM-EVs were administered and liver sections analysed by immunofluorescence and immunohistochemistry, using unlabelled AAM-EV–treated mice as controls. Biotin signal localised predominantly within and surrounding necrotic regions, consistent with uptake of AAM-EVs within injured tissue (Fig. S7C). Notably, biotin signal co-localised with enlarged LAMP1⁺ endolysosomal structures in necrotic areas, suggesting uptake through endocytic pathways and trafficking to the endolysosomal compartment (Fig. S7D).

To define the cellular distribution of AAM-EVs, NHS-AF647-labelled AAM-EVs were administered to mice with established APAP-ALI and liver analysed 1–6 h later (Fig. S8A– B). EV-associated fluorescence was rapidly detected in KCs and Mo/MoMFs (predominantly the Ly6Cint population), with comparatively little signal in neutrophils (Fig. S8B). Within the same livers, AF647+ monocytes were enriched for CD206 and showed markedly greater CD206 expression than their AF647− counterparts (Fig. S8C–D). This CD206-high phenotype was evident in AF647+ Ly6Clo cells from 1 h and maintained over 6 h, with a higher percentage of Ly6Chi cells becoming CD206 positive by 6h (Fig. S8E). Across the hepatic CD11b⁺ myeloid compartment, AAM-EV treatment was associated with increased CD206 positivity and expression within Ly6C^hi and Ly6C^int populations, but not KCs, by 3 h following administration (Fig. S8F), suggesting rapid phenotypic remodelling of recruited Mo/MoMFs.

### Extracellular vesicle release is required for AAM-mediated macrophage programming and therapeutic efficacy in liver injury

To determine whether EVs contribute to AAM function, EV release was inhibited using the neutral sphingomyelinase inhibitor GW4869. GW4869 treatment reduced EV secretion from cultured AAMs in a dose-dependent manner, decreasing particle concentrations by approximately 50%, 65% and 75–80% at 2.5 µM, 5 µM and 10 µM, respectively, with no change to cell number or viability (Fig. S9A). AAM phenotype was preserved at 2.5 µM and 5 µM, whereas 10 µM increased *Nos2* expression, indicating partial inflammatory activation. We therefore used 5 µM GW4869 for subsequent experiments. In real-time phagocytosis assays, GW4869 dose-dependently reduced the proportion of pHrodo⁺ AAMs and the intensity of particle uptake, indicating impaired efferocytic capacity when EV release was inhibited from AAMs (Fig. 5A). The reduced phagocytosis observed with GW4869-treated BMDMs could be partially rescued by treatment with AAM-EVs. We next examined the dependency of AAM efficacy on EV release *in vivo* using the APAP-ALI model (Fig. 5B). Whereas vehicle-treated AAMs significantly reduced hepatic necrosis and improved body weight recovery, GW4869-treated AAMs showed reduced therapeutic efficacy (Fig. 5C). Similarly, the enrichment of CD11b⁺TREM2⁺ macrophages within necrotic regions following AAM treatment was attenuated when EV release was inhibited. Across animals, the proportion of CD11b⁺TREM2⁺ macrophages inversely correlated with the extent of hepatic necrosis (Fig. 5D).

**Figure 5.**
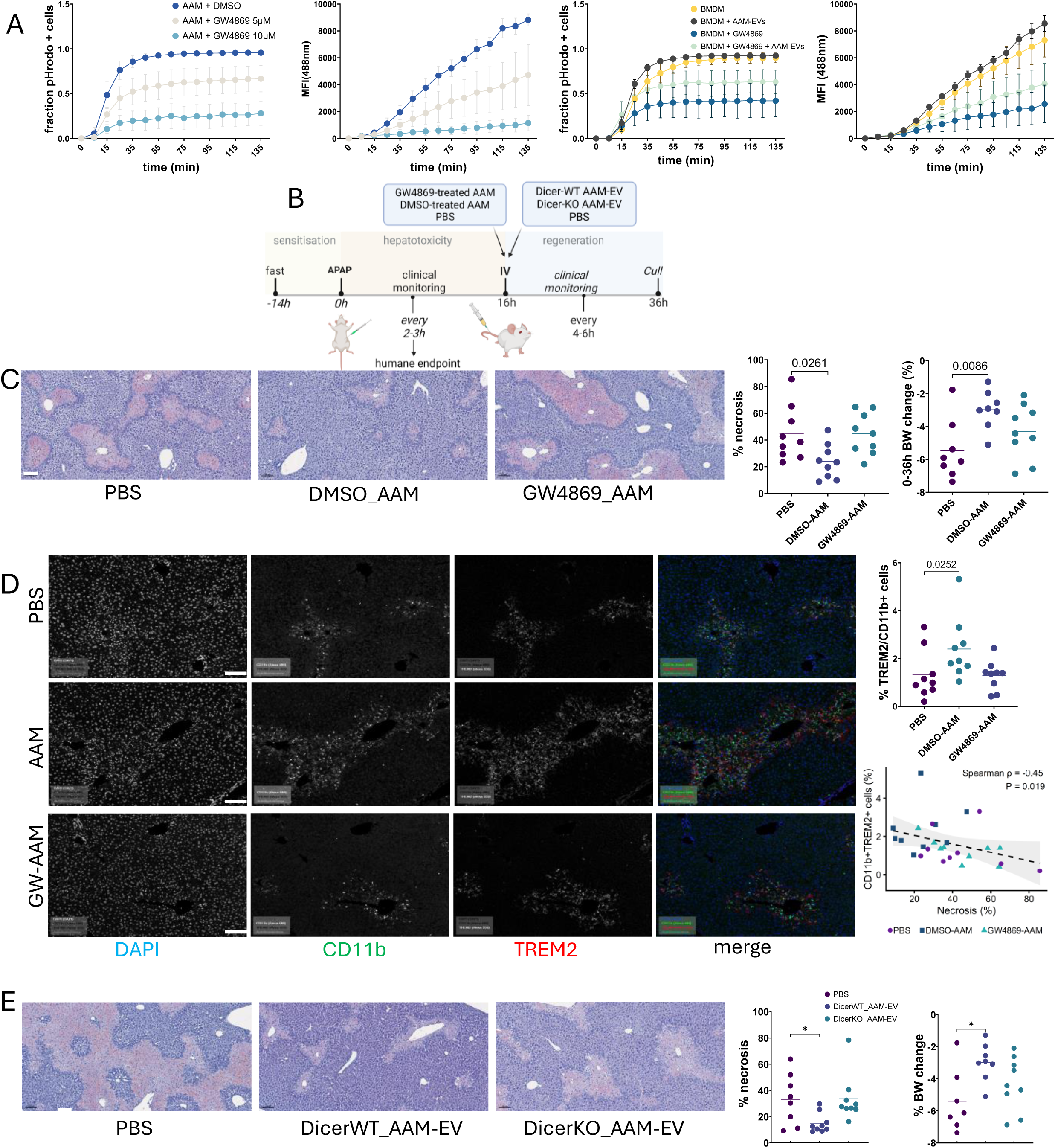
Extracellular vesicle release and microRNA processing contribute to AAM therapeutic efficacy. (A) Real-time phagocytosis assays showing the effects of GW4869 on AAM phagocytosis (left pair) and of AAM-EVs and/or GW4869 on BMDM phagocytosis (right pair). Within each graph pair, the fraction of pHrodo-positive cells and fluorescence intensity are shown over time. (B) Experimental schematic. APAP-ALI mice received AAMs (1 × 10⁶, i.v.) pre-treated with GW4869 or vehicle control, EVs derived from Dicer-WT or Dicer-KO AAMs, or PBS at 16 h post-injury, followed by clinical monitoring and tissue analysis at 36 h. (C) Representative HCE staining of liver sections from treated mice with quantification of hepatic necrosis and percentage body weight (%BW) change. Scale bars 100 µm. (D) Immunofluorescence staining of liver tissue showing CD11b (green), TREM2 (red), and DAPI (blue). Individual grayscale channels and merged images are shown, with quantification of CD11b⁺TREM2⁺ cells and correlation with hepatic necrosis (scale bars 100µm). (E) Representative histological images of liver injury in mice treated with EVs derived from Dicer-WT or Dicer-KO AAMs, with quantification of hepatic necrosis and percentage body weight change (right). Scale bars 50 µm.

### AAM-EV microRNA cargo contributes to therapeutic activity

EVs can mediate intercellular communication through the transfer of regulatory RNAs, including miRNAs, which can modify gene expression in recipient cells(*22*). To determine whether miRNA cargo contributes to AAM-EV therapeutic activity, *Dicer*^flox/flox bone marrow cells were treated with Tat-Cre recombinase to disrupt miRNA biogenesis prior to AAM differentiation. Dicer-deficient BMDMs retained the ability to upregulate expression of canonical AAM-associated markers following stimulation with IL-4 and IL-13 (Fig. S9B) and were used to generate DicerKO-AAM-EVs. Deletion of *Dicer* resulted in significantly reduced *Dicer* mRNA and undetectable expression of miRNAs including miR-125, miR-146 and miR-191 in EVs (Fig. S9B). In the APAP-ALI model (Fig. 5B), EVs derived from DicerKO-AAMs had no significant effect on necrosis, whereas EVs from DicerWT-AAMs were effective in reducing necrosis compared with PBS (Fig. 5E). As this suggested a functional contribution of EV-associated miRNAs to AAM-EV activity we sequenced the miRNA from AAM-EVs, BMDM-EVs and their parent cells. The miRNA composition of AAM-EVs was characterised using two complementary comparisons. Given enhanced efficacy of AAM-EV over BMDM-EV in functional assays, which reflected in differences in their parent cell efficacy in APAP-ALI(*4*), we compared the miRNA cargo of AAM-EVs with BMDM-EVs. This identified differences in EV miRNA representation associated with alternative macrophage activation, whereas comparison with parent AAMs assessed relative enrichment within the EV compartment. These comparisons identified partially overlapping miRNA populations (Fig. S9C). To examine the broader AAM-EV miRNA repertoire, we identified miRNAs showing both activation-associated and EV-relative enrichment. Among miRNAs detected in at least two of three AAM-EV samples with mean abundance ≥20 CPM, 25 showed >2-fold enrichment in AAM-EVs relative to both BMDM-EVs and parent AAMs (Fig. S9C). Separately, we considered four candidate miRNAs for target analysis. miR-149-5p, miR-451a and miR-27a-3p were more highly represented in AAM-EVs than BMDM-EVs (nominal P<0.05), while miR-181b-5p showed a similar directional difference (P=0.052). These candidates differed in their distribution between cells and EVs: miR-451a was strongly enriched in AAM-EVs relative to parent AAMs, whereas miR-149-5p, miR-27a-3p and miR-181b-5p were more highly represented in parent AAMs. Experimentally validated targets of these four candidate miRNAs included genes implicated in inflammatory signalling, DAMP/necrosis responses, leukocyte recruitment, cellular stress and anti-repair/fibrotic pathways in APAP-ALI (Fig. S9E). Functional enrichment of their validated targets identified pathways including endocytosis, ubiquitin-mediated proteolysis, protein processing, autophagy and mTOR signalling (Fig. S9F). Network analysis further demonstrated overlapping and distinct validated interactions between the four candidate miRNAs and APAP-ALI-associated genes (Fig. S9G).

### AAMs and AAM-EVs induce overlapping but distinct transcriptional responses in the injured hepatic niche

To compare the effects of AAMs and their extracellular vesicles, we generated GFP-labelled AAMs and AAM-EVs, both of which retained therapeutic activity (Fig. S10). Single-nucleus RNA sequencing was performed on livers from mice treated with PBS, AAMs or AAM-EVs at 16 h after APAP administration and collected at 36 h (Fig. 6A). GFP-positive cells were observed within and around necrotic regions following AAM administration, whereas GFP signal was not detected following AAM-EV treatment. Reclustering of the myeloid compartment identified Kupffer cells (KCs), monocytes/monocyte-derived macrophages (MoMFs), dendritic cells and a small neutrophil population (Fig. 6B; Fig. S11A).

**FIGURE 6.**
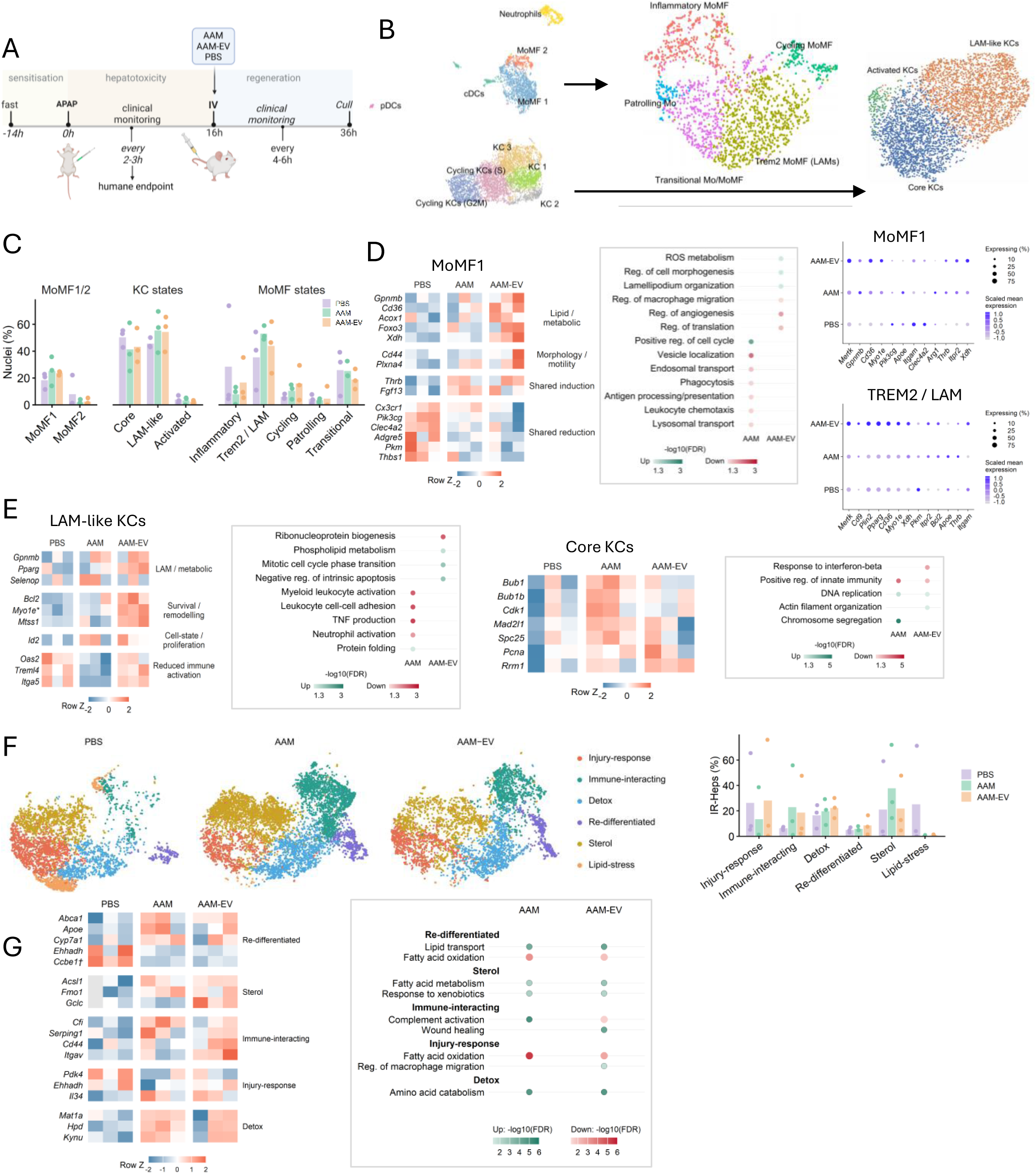
AAMs and AAM-EVs induce convergent remodelling of the injured hepatic niche in APAP-ALI. (A) Experimental design for snRNA-seq of APAP-injured livers treated with PBS, AAMs or AAM-EVs. (B) UMAPs showing the myeloid compartment and reclustering of MoMFs and KCs into the indicated transcriptional states. (C) Relative abundance of MoMF1/2, KC states and MoMF states by treatment. Bars represent mean animal-level proportions and dots represent individual animals. Proportions are calculated within the corresponding parent population. (D) Animal-level expression heatmap and representative GO Biological Process enrichment for treatment-associated changes in MoMF1, with dotplots showing selected treatment-associated genes in the indicated MoMF populations. Dot size indicates the percentage of nuclei expressing each gene and colour indicates scaled mean expression. (E) Animal-level expression heatmaps and representative GO Biological Process enrichment for treatment-associated changes in LAM-like and core KCs. (F) Condition-split UMAPs showing six IR-Hep states, with their relative abundance by treatment. Bars represent mean animal-level proportions and dots represent individual animals. (G) Animal-level expression heatmap of selected treatment-associated genes, grouped by IR-Hep state, with corresponding functional enrichment summaries. Heatmap colours indicate row-scaled normalised expression. In pathway plots, green and red indicate enrichment among increased and decreased genes, respectively, with colour intensity indicating −log₁₀(pathway FDR). Treatment comparisons are against PBS.

Within MoMF1, AAM and AAM-EV treatment were associated with partially overlapping transcriptional responses, including increased Thrb, *Fgf13* and *Plxna4* and reduced *Thbs1*, *Pik3cg*, *Clec4a2* and *Adgre5*. AAM-EVs were additionally associated with increased expression of lipid- and oxidative-metabolism-associated genes, including *Gpnmb*, *Cd36*, *Acox1* and *Xdh*. Pathway analysis identified reactive oxygen species metabolism and cell morphogenesis among AAM-EV-associated programmes, whereas AAM treatment was associated with cell-cycle-related pathways and reduced representation of phagocytosis, antigen presentation and leukocyte chemotaxis pathways (Fig. 6D).

Further reclustering of MoMFs resolved five states: inflammatory, *Trem2*/LAM-like, cycling, patrolling monocytes and transitional Mo/MoMFs (Fig. 6B; Fig. S11B). Both treated groups showed a higher mean proportion of *Trem2*/LAM-like macrophages than PBS, although this varied between animals (Fig. 6C). Complementary exploratory MAST analysis of pooled nuclei identified treatment-associated candidates involved in phagocytosis, lipid handling and cytoskeletal organisation, including *Mertk*, *Cd36*, *Pparg*, *Myo1e* and *Mtss1*, together with reduced expression of inflammatory-associated genes such as *Pik3cg* and *Il1b* (Fig. 6D). These findings supported further examination of macrophage programmes associated with clearance and tissue remodelling.

KC reclustering identified core, LAM-like and a smaller activated population (Fig. 6B–C; Fig. S11B). Within LAM-like KCs, both treatments altered lipid-handling and reparative-state genes, including *Gpnmb*, *Pparg* and *Selenop*. AAM-EV treatment was associated with a membrane–cytoskeletal remodelling signature characterised by increased *Myo1e*, *Mtss1* and *Sh3pxd2b*, alongside *Gpnmb* upregulation. Notably, *Myo1e*, an unconventional myosin implicated in phagocytic cup formation, was significantly increased following AAM-EV treatment (pseudobulk FDR = 0.017). Pathway analysis identified reduced representation of TNF production, myeloid activation and leukocyte adhesion pathways following AAM treatment, whereas AAM-EVs were associated with phospholipid metabolism and mitotic cell-cycle phase transition among upregulated genes, and ribonucleoprotein complex biogenesis among downregulated genes (Fig. 6E). Together, these analyses identify overlapping macrophage responses to AAMs and AAM-EVs, alongside exploratory differences in metabolic, cytoskeletal and cell-cycle-associated programmes. These differences may reflect distinct properties of parent cells and isolated EVs, although differences in dose, tissue distribution and exposure kinetics may also contribute.

In core KCs, AAM treatment increased expression of cell-cycle-associated genes, including *Bub1*, *Bub1b*, *Cdk1* and *Pcna*, with enrichment of chromosome segregation and DNA replication pathways. AAM-EV-associated changes included enrichment of DNA replication and actin filament organisation among upregulated genes, and reduced representation of interferon and innate immune-response pathways (Fig. 6E).

To further explore the changes in cycling programmes, we conducted MAST analysis on pooled nuclei in different KC states by condition and observed a broad cycling-associated response to AAM treatment, whereas the AAM-EV cycling response appeared to be concentrated in the small population of activated KCs (Fig. S11D).

Together, these analyses identify overlapping macrophage responses to AAMs and AAM-EVs, with differences in metabolic, cytoskeletal and cell-cycle-associated programmes.

### Hepatocyte compartment

Reclustering 14,996 *Anxa2*⁺ IR-Hep nuclei identified six states encompassing injury-response, immune-interacting, detoxification, re-differentiated, sterol and lipid-stress programmes (Fig. 6F). The injury-response cluster expressed stress-associated genes including *Plat* and *Gdf15* and was enriched for stress response, IL-1 and cell motility pathways. Immune-interacting hepatocytes expressed cytokine- and complement-associated genes, including *Il1r1* and *Cfh*, with enrichment of immune communication and migratory programmes. Additional states reflected metabolic adaptation, including detoxification hepatocytes enriched for xenobiotic and glutathione metabolism pathways, re-differentiated hepatocytes expressing mature metabolic genes including *Mat1a* and *Cps1*, sterol-associated hepatocytes enriched for cholesterol biosynthesis pathways, and lipid-stress hepatocytes characterised by fatty acid degradation and PPAR signalling programmes (Fig. 6F; Fig. S12A). State composition varied between animals (Fig. 6F), but we found that lipid-stress and injury-response states were associated with higher ALT and necrosis and greater body weight loss, whereas sterol, re-differentiated and immune-interacting states showed the opposite pattern (Fig. S12B).

State-specific differential expression and pathway analysis identified both shared and distinct responses to AAM and AAM-EV treatment. At the whole-compartment level, treatment-associated fold changes were positively correlated (Spearman ρ = 0.76; Fig. S12B). In re-differentiated IR-Heps, both treatments were associated with increased expression of lipid-handling gene candidates including *Abca1*, *Apoe* and *Cyp7a1* and reduced *Ehhadh*, alongside enrichment of lipid transport and homeostasis among upregulated genes and fatty acid oxidation among downregulated genes. Sterol-associated hepatocytes showed increased *Acsl1* and *Fmo1*, consistent with enrichment of fatty acid metabolism and responses to xenobiotics. In detoxification hepatocytes, increased *Mat1a*, *Hpd* and *Kynu* accompanied enrichment of amino acid catabolism. Immune-interacting hepatocytes showed increased complement-regulatory genes, including *Cfi* and *Serping1*, following AAM treatment, whereas AAM-EV treatment increased adhesion-associated genes *Cd44* and *Itgav*, alongside wound-healing enrichment. Injury-response hepatocytes showed reduced *Pdk4* and *Ehhadh* following both treatments, while AAM-EV treatment additionally increased *Il34*, providing a candidate link to macrophage signalling (Fig. 6G). These findings suggest partially shared metabolic responses to treatment, alongside distinct changes in hepatocyte immune communication and tissue remodelling.

We also observed close apposition between TREM2⁺ macrophages and ANXA2⁺ hepatocytes at the edge of pericentral regions in APAP-injured tissue (Fig. S12C). CellChat predicted IR-Hep-to-TREM2 macrophage signalling through Pros1–Mertk, Apoa1–Abca1, Lamc1–Cd44 and Lgals9–Cd45 across PBS, AAM and AAM-EV conditions, with Fn1–Cd44 additionally predicted in PBS and AAM-EV conditions (Fig. S12D). These predictions provide potential molecular links between the adjacent hepatocyte and macrophage populations. Our analysis extends the characterisation of Anxa2⁺ injury-responsive hepatocytes by resolving six transcriptionally distinct states following APAP injury and identifying their responses to AAM and AAM-EV treatment.

## DISCUSSION

This study identifies extracellular vesicle-mediated communication as a previously unrecognised mechanism contributing to the therapeutic efficacy of AAMs in acute liver injury. AAM-derived EVs enhanced macrophage phagocytosis and promoted reparative gene expression *in vitro*, and recapitulated the effects of their parent cells *in vivo*, reducing hepatic necrosis and neutrophilic inflammation and enhancing hepatocellular regeneration *in vivo*. Pharmacological inhibition of EV release attenuated the therapeutic efficacy of parent AAMs, while EVs derived from Dicer-deficient macrophages showed reduced therapeutic activity. Together, these findings implicate EV secretion and miRNA-associated mechanisms in AAM-mediated liver repair and support the potential therapeutic development of AAM-EVs.

We observed remodelling of endogenous macrophage populations within the injured liver following human AAM therapy, with increased TREM2⁺ macrophages in necrotic regions, altered *Gpnmb* expression and downregulation of inflammatory myeloid programmes. TREM2 has an established role in hepatic injury resolution, including modulation of TLR/NF-κB signalling, phagocytosis and the acquisition of reparative macrophage phenotypes(*14, 15*). More recently, De Ponti and colleagues demonstrated that ontogenically distinct recruited macrophages and LAM-like Kupffer cells exhibit complementary contributions during liver repair, with TREM2-dependent efferocytosis contributing to their reparative activity. They further identified efferocytosis within pericentral injury regions as a driver of LAM-like macrophage identity(*23*). Consistent with these observations, AAM-EVs enhanced macrophage phagocytosis *in vitro*, while proteomic analysis of recipient macrophages identified increased TREM2 abundance and enrichment of endocytosis, Fc receptor-mediated phagocytosis and intracellular signalling pathways. The increased representation of TREM2/GPNMB-associated macrophage programmes following AAM and AAM-EV treatment may therefore reflect enhanced clearance of damaged cells and acquisition of reparative macrophage states. Given that transcriptional profiling was performed 20 h after treatment, when hepatic necrosis and inflammation were already reduced, this may reflect a later phase of macrophage-mediated resolution following earlier phagocytic activity. Together, these findings suggest that EV-mediated communication contributes to macrophage reprogramming during liver repair.

Our biodistribution studies identify hepatic macrophages and monocytes as prominent recipients of AAM-EVs, with preferential localisation to injured regions and LAMP1⁺ endolysosomal compartments. Importantly, EV association was accompanied by early changes in myeloid phenotype, including increasing CD206 expression in hepatic monocyte populations within hours of administration. These findings raise the possibility that AAM-EVs provide a form of phenotypic instruction, transmitting aspects of the parent macrophage’s reparative programme to endogenous myeloid cells. In line with this, a number of studies have demonstrated that macrophage EVs can alter naïve macrophage gene expression to acquire phenotypic characteristics of the parent cell, and can also alter circulating inflammatory monocyte populations (*19, 24*). Together, the spatial distribution and early phenotypic responses support a model in which AAM-EVs communicate with endogenous myeloid cells to promote a reparative hepatic environment.

Comparative transcriptomic analyses revealed overlapping but distinct responses to AAMs and AAM-EVs. Both treatments were associated with changes in macrophage lipid metabolism, inflammatory signalling and cellular remodelling. Notably, *Myo1e* was increased in LAM-like KCs following AAM-EV treatment, with a similar increase observed in *Trem2* macrophages following hAAM therapy in an independent sequencing study. This protein is implicated in actin-dependent membrane dynamics and phagocytic cup formation, providing a potential mechanistic link between the transcriptional response to AAM-EVs and the enhanced phagocytic activity observed *in vitro*(*25*). Additional treatment-associated changes in cytoskeletal genes, including *Mtss1* and *Sh3pxd2b*, suggest that membrane–actin remodelling may contribute to the macrophage response to AAM-EVs. The differences between parent AAMs and AAM-EVs may also provide insight into their mechanisms of action. Although both treatments altered macrophage inflammatory and metabolic programmes, AAM-EVs were more clearly associated with cytoskeletal and lipid-handling responses in LAM-like KCs, whereas parent AAMs showed greater enrichment of cell-cycle-associated programmes. These differences (at 20 hours after administration) may reflect the capacity of living macrophages to respond dynamically to environmental signals and their continued release of AAM-EVs, compared with the delivery of a finite molecular cargo by EVs. Nevertheless the reductions in necrosis were remarkably similar, despite the probable differences in dose, exposure kinetics and spatial location.

Beyond macrophages, hAAM therapy was associated with substantial remodelling of hepatocyte states, characterised by enrichment of metabolic and biosynthetic programmes, including fatty-acid synthesis, alongside reduced representation of lipid catabolic pathways. These findings suggest that macrophage therapy influences not only inflammatory resolution but also the restoration of parenchymal function. Comparative transcriptomics revealed partially overlapping hepatocyte responses to AAMs and AAM-EVs, consistent with their shared therapeutic effects. Whether these changes reflect direct EV signalling to hepatocytes, altered macrophage-derived signals or secondary consequences of reduced injury remains unresolved. Temporal analysis will be important to distinguish these mechanisms.

Our mechanistic perturbation experiments implicate EV secretion and miRNA-associated cargo in AAM therapeutic activity. The reduced efficacy of GW4869-treated AAMs, together with attenuated enrichment of TREM2⁺ macrophages within necrotic regions supports a role for EV mediated communication in the reparative effects of parent macrophages. Although GW4869 may affect sphingolipid metabolism independently of EV secretion, the concentration used did not alter the measured phenotype of parent AAMs, supporting the interpretation that reduced EV release contributed to the loss of therapeutic efficacy. Moreover, reduced efficacy of Dicer-deficient AAM-EVs supports a contribution from miRNA-associated mechanisms, while any residual therapeutic activity may reflect contributions from Dicer-independent miRNAs, such as miR-451a, or EV-associated proteins and/or lipids. Depletion of miR cargo using the Dicer model has also been shown to prevent the improved glucose tolerance and insulin sensitivity observed in obese mice administered macrophage EVs(*26*). Notably, our unbiased miRNA sequencing identified candidate miRNAs, including miR-27a-3p, which has previously been shown to promote CD206 expression and enhance phagocytic activity in recipient monocytes, and has also been implicated in the therapeutic activity of EVs from MSCs (*27, 28*). Another miR upregulated in AAM-EVs, miR-181b-5p has been implicated in M2-EV mediated cardiac repair(*29*). However, there is some evidence that EV-associated proteins may also contribute to therapeutic activity, as heat treatment has been shown to impact the efficacy of macrophage derived EVs (*30*).

These findings have important translational implications. AAM-EVs reproduced several beneficial effects of parent AAMs, with a significant dose-dependent reduction in hepatic necrosis. Indeed, previous studies have demonstrated that EVs from macrophages can suppress inflammation and modulate recipient myeloid cells in models of atherosclerosis, metabolic and cardiac disease, albeit with higher particle doses and repeated administrations (*19, 24, 31*). Importantly, AAM-EVs retained therapeutic efficacy when administered during established APAP injury at a dose approximating the 24h output of the efficacious parent-cell dose. However, EV dose does not necessarily correlate linearly with biological activity since previous *in vitro* cell stimulation work showed that lower EV doses elicited more cell-source-specific transcriptional responses, whereas higher doses were associated with changes in lysosomal and membrane-trafficking programmes, suggesting that the cell becomes occupied with EV uptake (*17*). Many studies examining therapeutic EVs, particularly those from mesenchymal stem cells (MSCs), have observed efficacy when the EVs are administered alongside or shortly after the hepatotoxic insult(*32, 33*). In contrast, the delayed therapeutic window in our study supports the potential relevance of AAM-EVs to patients presenting after hepatic injury has developed, when N-acetylcysteine is less effective(*21*). As a cell-free product derived from a well-defined parent cell population, AAM-EVs could potentially offer advantages in standardisation, safety and storage. Translation will require confirmation using clinically manufactured human AAM-EVs, reproducible production and potency assays, and investigation of biodistribution, therapeutic dosing and efficacy across a wider range of injury phenotypes and timepoints.

Together, these findings identify EV-mediated communication as an important mechanism through which therapeutic macrophages influence endogenous myeloid responses and promote liver repair. By linking EV secretion, miRNA-associated cargo and macrophage phenotypic remodelling to therapeutic efficacy, this study advances our understanding of macrophage-mediated tissue repair and provides a foundation for developing cell-free macrophage-derived therapeutics.

## Acknowledgements

We thank Fiona Rossi and members of the IRR Flow Cytometry facility for assistance with flow cytometry and FACS. We thank Richard Clarke, Angie Fawkes and staff of the Edinburgh Clinical Research Facility for assistance with RNA sequencing. We thank Meryam Beniazza for assistance with 10x single nucleus sequencing, Dr Rob Hunter for assistance with EV isolation techniques and Kirsten Wilson for liver enzyme assays. We thank all the staff at the IRR Imaging Facility, University of Edinburgh for assistance with high content imaging and confocal microscopy. This work has made use of the resources provided by the Edinburgh Compute and Data Facility (ECDF). For the purpose of open access, the author has applied a Creative Commons Attribution (CC BY) licence to any Author Accepted Manuscript version arising from this submission. This work was funded by the Medical Research Council (MRC) (Funding Reference No. MR/M007588/1) and the Chief Scientist Office grant “Precision Cell Therapy for the Liver”.

## Methods & Materials

### BMDM and AAM production

Mouse bone marrow (BM) suspension was flushed from femurs and tibias of healthy C57/BL6 mice (8-12 weeks old, Charles River) or age-matched MacGreen mice (expressing Green fluorescent protein under the *Csf1r* promoter), filtered (70 µm), and incubated into ultra-low attachment flasks (Corning Inc.) in presence of DMEM:F12 (1:1) cell culture media (Gibco) supplemented with FBS (10%), L-glutamine (2 mM), penicillin/streptomycin (100 U/ml, 100 μg/ml), and murine recombinant CSF1 (40 ng/ml; Peprotech) for 6 days (37 C, 5% CO2), with additional feeds on day 3 and 5 (20 ng/ml CSF1, in a 50% media change). To produce AAMs, BMDMs from all mice were combined, counted and resuspended in flasks with medium containing interleukin (IL)-4 and IL-13 (Miltenyi); 20 ng/ml each overnight. Cells for injection were frozen in cryostor (CS-10) and stored in liquid nitrogen until the day of administration.

### hAAM production

CD14+ monocytes were isolated and cultured as previously described(*4*). Briefly, they were isolated from a UK donor leukapheresis product using the CliniMACS Prodigy® closed system, cultured for 6 days (37 °C, 5% CO2) in low adhesion culture bags (Miltenyi) in the presence of TexMACSTM serum-free media (Miltenyi) and 24,600 IU/ml CSF1 (100 ng/ml, RCD Systems). For hAAM polarization cytokines IL-4 and IL-13 were added on day six of culture at 400 IU/ml and 60 IU/ml, respectively. On day 7, hAAM were harvested, washed and formulated at the required final cell density in PBS and 66% CryoStor cell cryopreservation media (CS10 Medium) and filled into CryoMACS bags and vials. Successful hMDM differentiation was confirmed using flow cytometry to demonstrate a ≥5-fold increase in MFI on CD206 and 25F9 expression compared to initial CD14+ cells from day 0. Cell viability was performed using Trypan Blue staining and Burker Haemocytometer Counting Chambers before the injection, and the cells were maintained at 4 °C until the time of injection (∼10 min).

### APAP-ALI and treatment administration

Ten to twelve week-old male C57BL6/J mice, averaging 25-30g (Charles River, UK) housed in open top cages were fasted for 14 h. All mice received a single APAP injection (350mg/kg, i.p., Sigma Aldrich) in warm sterile saline (PanReac Applichem) or saline only for uninjured controls. Standard chow, wet mash and enrichment bedding/housing were returned to mice 20 min post-APAP administration. Thawed human or mouse AAMs (1–5 × 10^6^ cells, i.v., 100–200 μl) or AAM-EVs (1×10^6^ – 4×10^8^ particles, i.v., 100µl) were resuspended in PBS (Sigma) for mouse AAMs, or PBS and administered to APAP-ALI mice at 16 h after APAP injury. For EV experiments, control preparations consisted of unconditioned culture medium processed through the same ultrafiltration and size-exclusion chromatography steps as conditioned medium. These preparations are labelled ‘PBS’ throughout the manuscript and figures. Following i.v. injections all mice were transferred to a warming cabinet (28°C). Approximately 1 hour before cull, mice were pulsed with 5-bromo-2′-deoxyuridine (BRDU, 1 mg in sterile saline, i.p., Sigma Aldrich) to label proliferating cells. The mice were anaesthetized with isoflurane for cardiac exsanguination to collect blood, then culled via cervical dislocation; death was confirmed by exsanguination through severing the femoral artery. Liver was perfused with 10ml PBS and fixed in 10% of formalin for 8 hours. All animal experiments were conducted in accordance with the criteria outlined in a license granted under the Animals (Scientific Procedures) Act 1986 and were approved by the University of Edinburgh Animal Ethics Committee.

### Imaging/biodistribution studies

Prior to baseline scans mice were depilated to remove all fur covering the lungs and abdomen. For all scans mice were anaesthetised with 3% (v/v) isoflurane/O2 in a pre-heated chamber, then transferred to the pre-heated scanning bed of the Biospace PhotonIMAGER OPTIMA (Biospace Lab, France) where they were maintained under 2% (v/v) isoflurane/O2 for imaging. Mice were positioned prone on the scanner and x-rays and near-infrared fluorescence images taken using Biospace Photon Acquisition software V3.2. Fluorescence imaging used a 50mm f/1.2 Lens with 1.2 Aperture and excitation/emission settings of 737/797 nm with a 780-815 nm band pass filter and exposure of 1000ms per frame with 5 frames taken and averaged to produce a composite scan image for each mouse/timepoint. Autofluorescence background of 437 nm was automatically subtracted prior to imaging analysis. Fluorescence images were co-registered with x-rays and in vivo near-infrared signal in the liver region was analysed using M3 Vision software. To confirm in vivo imaging results, following sacrifice, mice were dissected and organs imaged ex vivo using the same excitation/emission settings with brightfield co-registration. Results for in vivo images/organs are reported in photons/second/cm2/steradian to account for the imaged area and 3D nature of the mouse/tissue. Near-infrared labelling of the EVs was used to enable sufficient fluorescence signal penetration from deep tissues whilst avoiding absorption from skin, lipids and haemoglobin.

### Liver digestion and flow cytometry

Hepatic non-parenchymal cells (NPCs) were isolated following perfusion of murine livers through the inferior vena cava with 10 mL phosphate-buffered saline (PBS). Livers were excised and weighed, and approximately 0.5 g of tissue was finely minced and digested in serum-free RPMI containing collagenase V (0.8 mg/mL; Sigma-Aldrich), collagenase D (0.625 mg/mL; Roche), dispase (1 mg/mL; Gibco) and DNase I (30 µg/mL; Roche) at 37°C with shaking (240 rpm) for 15–25 min. Digested tissue was passed through a 100 µm cell strainer, washed and centrifuged at 300 × g for 5 min at 4°C. Following erythrocyte lysis, cells were washed, filtered through a 40 µm strainer and resuspended in FACS buffer (PBS containing 0.5% fetal bovine serum and 2 mM EDTA). For phenotypic characterisation of cultured AAMs and BMDMs, cells were harvested and stained with antibodies targeting macrophage surface markers. Non-specific antibody binding was blocked using 10% BSA for 15 min at 4°C, followed by incubation with fluorophore-conjugated antibodies (1:200 dilution) for 30 min at 4°C in the dark. For hepatic NPC analysis, cells were counted and stained with LIVE/DEAD Fixable Near-IR viability dye (Invitrogen) for 30 min on ice, followed by Fc receptor blocking with anti-CD16/32 (BioLegend; 1:200) for 10 min. Cells were subsequently incubated with fluorophore-conjugated antibodies (1:200 dilution) in True-Stain Multi-Fluor Buffer (BioLegend) for 30 min at 4°C in the dark. Following staining, hepatic NPCs were washed, fixed in 4% paraformaldehyde for 15 min at room temperature and resuspended in FACS buffer. Antibody panels were selected to distinguish macrophage populations and activation phenotypes, including Kupffer cells, monocytes/monocyte-derived macrophages and neutrophils (Supplementary Table 1). Unstained, fluorescence-minus-one and single-stained compensation controls were included as appropriate. Samples were acquired using a five-laser BD LSRFortessa flow cytometer (BD Biosciences, USA). Gating strategies are provided in the corresponding supplementary figures.

### Immunostaining

Four-micron sections of formalin-fixed paraffin-embedded (FFPE) block slides were dewaxed, rehydrated, stained with HCE, and mounted with Pertex mounting medium (Histolab). For necrosis quantification, HCE sections were scanned on a Vectra Polaris Automated Quantitative Imaging System (PerkinElmer) using multispectral imaging. At least six/seven 10x ROIs (representing a minimum combined tissue area of 15.5 mm2) were analyzed for each sample with InForm automated image analysis software (AKOYA Biosciences), which allows to design an algorithm based on the presence of the healthy, necrotic, and previous damaged tissue. After training using a representative image from every sample, and then batch processed on all images, necrosis was quantified by expressing necrotic tissue area as a percentage of total tissue area.

### Immunodetection assays and quantification

Four-micron sections of FFPE blocks slides were dewaxed and rehydrated before heat-induced antigen retrieval. After blocking in protein solution (Spring Bioscience) for 30 min, the slides were incubated overnight with primary antibodies at 4 °C (see supplementary table): Anti-Ly6G Biolegend, Anti-Ki67 Abcam, Anti-BRDU or isotype-control IgG (Vector Laboratories). For BRDU staining, successful BRDU injection was confirmed first by examining BRDU in highly proliferative tissue such as intestine (animals were removed from the analysis if no staining was present here). For immunofluorescence, the slides were incubated with secondary antibodies for 30 min in the dark: Alexa Fluor® 488, 647 conjugate, and DAPI (SIGMA ALDRICH, 1 µg/ml, 10 min) was applied before mounting in Fluoromount-G (Invitrogen). The intensity of the fluorescent signal is quantified in DAPI-positive nuclei. For quantification, ten fields per tissue were analyzed at Polaris multispectral analysis (PerkinElmer) and InForm (AKOYA Biosciences) automated image analysis software. After segmentation of the tissues and the scoring of all the images based on the positivity threshold, an algorithm created on the presence of the double positive cells was designed, expressing the percentage of the total cells number. For immunohistochemical staining, a biotinylated secondary antibody was incubated (1:500) for 1 h before treatment with an avidin-based peroxidase reagent (Vector Laboratories). After DAB (3, 3’-diaminobenzidine) colorimetric development using a Harris’ hematoxylin counterstain, the sections were washed in acid alcohol and Scott’s tap water substitute, dehydrated, and mounted with ClearVue mountant (ThermoScientific). For quantification, after Polaris multispectral analysis (PerkinElmer), ten 10x ROIs per tissue were analyzed with InForm (AKOYA Biosciences) automated image analysis software which allows designing an algorithm based on the presence of the positive cells expressed as a percentage on the total cells number (hematoxylin staining was used for the counterstaining).

### TUNEL staining

For in situ cell death detection, four micron sections of FFPE block slides were dewaxed and rehydrated before proteinase K antigen retrieval. Slides were incubated with TUNEL reaction mixture (Roche) for 20 min at 37 °C in the dark, following manufacturer’s instruction. After DAPI (SIGMA ALDRICH, 1 µg/ml, 10 min) incubation, slides were mounted in Fluoromount-G (Invitrogen) and analyzed under a fluorescence microscope Nikon Eclipse Ni.

### Serum analyses

Serum was isolated from whole blood after centrifugations at 1500 × g and 14,000 × g for 5 min each. Serum chemistry was performed by measurement of ALP, ALT, AST, creatinine and albumin utilizing a commercial kit (Alpha Laboratories Ltd). All kits were adapted for use on a Cobas Fara centrifugal analyzer (Roche Diagnostics Ltd). For all assays, intra-run precision was CV < 4%. Cytokines (TNF-α, IFN-γ, IL-1β, IL-2, IL-5, KG/GRO, IL-6, IL-10) were quantified in serum using a V-plex Pro-inflammatory Panel kit (Meso Scale Discovery) following the manufacturer’s instructions. The plate was read on a QuickPlex SQ 120 analyzer (Meso Scale Discovery). Standards were assayed in duplicate and samples in singlet as recommended.

### EV harvesting, isolation and quality control

BMDMs were plated at a density of 5 million per 150mm round tissue culture treated plate and exposed to 20ng/ml mCSF, IL-4 and IL-13 overnight to generate AAMs (or mCSF only to maintain BMDMs). The following day medium was removed and cells were washed twice with warmed PBS, and cultured with complete medium containing 10% reduced bovine serum FCS (Miltenyi, EXO-FBS-50A-1) and 20ng/ml IL-4 and IL-13 (Miltenyi) to maintain AAM phenotype. EVs were harvested using a method based on differential ultracentrifugation, ultrafiltration and size exclusion chromatography (SEC)(*18*). Conditioned culture media (CCM) was harvested from an average of 20 plates with 5×10^6^ cells/plate every 24h until 2 harvests were done. RNA was sampled from AAM or BMDM at each time point to measure polarisation markers. CCM was subjected to serial spins of 300g (15 minutes) to remove cells and 5000g (15 minutes) to remove debris and was then filtered through 0.2µm mesh to remove microvesicles. The filtered supernatant was then loaded into 15ml 100kDa Amicon filters (UFC901024D) with 15ml CCM and spun at 4000g until concentrate to approximately 500µl. Concentrated CCM was then stained with DiIC18(7) (1,1’-Dioctadecyl-3,3,3’,3’-Tetramethylindotricarbocyanine Iodide) (DiR) by incubating with 5µM DiR in the dark at room temperature for 30 minutes, or with 2µM PKH-26 (Mini26-kit, Merck) in the dark for 10 minutes. To separate EVs from proteins and free dye, 2 x 500µl CCM was serially loaded onto Exospin Midi size exclusion columns (Cell Guidance Systems), and 24 fractions of 500µl were collected, using PBS as the eluent. Protein concentrations of fractions were determined using a Bradford assay (Bio-rad). Particle concentration of fractions was determined with nanoparticle tracking analysis using a Zetaview (Analytic). As particle:protein ratio revealed the EV rich fractions to be 7-12, with protein and dye eluting in the later fractions, these were pooled and reconcentrated using a 10kDa Amicon filter. Medium (containing 10% reduced bovine serum FCS) unexposed to cells was subject to the same procedure as the CCM (30ml concentrated to 1ml using UF and run through SEC column with fractions 7–12 pooled, re-concentrated using UF and subject to NTA analysis) and was used as the control in all experiments and is referred to as ‘PBS’ for clarity in graphs. For staining of EVs, the CCM (or the same media unexposed to cells) was exposed to NHS Ester 647 (A37573, Thermo Fisher) or PKH-26 (Merck) and labelled according to kit instructions. Dye was then washed off using SEC, with confirmation of free dye eluting at later fractions, with further concentration of pooled fractions using a 10kDa Amicon filter. For studies using 647 or PKH-26 labeled EVs, the appropriate wash control was administered to account for any signal relating to free dye.

For transmission electron microscopy, 5µl of concentrated EV samples were adsorbed onto formvar-coated copper grids and negatively stained with 2% uranyl acetate for contrast. Grids were allowed to air dry prior to imaging. TEM was performed using a JEOL-1400 transmission electron microscope operating at 80 kV, equipped with a Gatan OneView digital camera. Representative images were acquired at appropriate magnifications, with settings optimized for EV visualization.

For imaging of EV uptake, BMDMs were plated onto Ibidi γ 8 well plates at a density of 5×10^4^ cells/well and were imaged at 63x water objective (NA 1.1) in the orange\red channel (561nm) and Brightfield on an Opera Phenix™ Plus (Revvity).

### Western blotting

For western blotting, EVs were lysed with RIPA (Thermofisher) supplemented with 1:100 protease inhibitor cocktail (Halt) and sonicated for 30 seconds. Parent cells were lysed in 100ul/million cells RIPA supplemented with 1:100 protease inhibitor cocktail and then agitated for 30 mins at 4C, followed by a 15,000g spin for 20 min at 4C before subjecting the supernatant to Bradford assay. Cell or EV sample was then heated to 70C for 10 minutes and loaded onto an increased capacity Bolt 4-12% 1mm mini protein gel (Thermofisher), and electrophoretically transferred to nitrocellulose membranes (Invitrogen). The membranes were then blocked with 5% bovine serum albumin and incubated with primary antibodies (see supplementary table 1) with constant gentle agitation overnight, followed by 3 washes with TBST. The membranes were then incubated with respective HRP-conjugated secondary antibodies (1:5000) and chemiluminescence was captured on a ChemiDocTM MP Imaging System (Bio-Rad).

### Real Time Phagocytosis Assay

BMDMs were plated on 96 well plate (Revvity, CellCarrier) at a density of 1×10^5^ and allowed to attach. They were pretreated overnight with PBS, BMDM-EVs or AAM-EVs at a 100:1 EV:cell ratio. Cells were stained with CellMask Deep Red (Thermo Fisher) and NucBlue (Thermo Fisher) and washed with PBS. Phagocytosis was initiated by the addition of pHrodo™ green Zymosan A particles (ThermoFisher). Fluorescent images were taken at Opera Phenix™ Plus (Revvity) using 20x water objective (NA 1.0) in the 4′,6-diamidino-2-phenylindole (DAPI) channel, 488 nm, and 647 nm before, and at 5 min intervals after the addition of bioparticles for a maximum of 150 min. Macrophages positive for phagocytosis were classified based on mean fluorescence intensity (488 nm) greater than 500 and expressed as pHrodo positive fraction of all live cells (NucBlue positive cells). Phagocytosis measurements were obtained from 6–8 wells per group, with repeated measurements from the same wells over time.

### Proteomics

Cell and EV samples were processed at the University of Edinburgh Proteomics Core. Proteins were extracted, reduced, alkylated and digested with trypsin prior to C18 reverse-phase peptide clean-up and Orbitrap mass spectrometry analysis. Raw data were processed using MaxQuant. Recipient-cell label-free quantification (LFQ) data were analysed in R using the DEP workflow. Protein groups quantified in all three replicates of at least one treatment group were retained. Intensities underwent variance-stabilising normalisation, and missing values were imputed using the probabilistic minimum method (MinProb; q = 0.01). Differential protein abundance was assessed using limma linear models with empirical Bayes moderation, comparing AAM-EV-treated and BMDM-EV-treated macrophages with PBS-treated macrophages and directly comparing the two EV treatments. Benjamini–Hochberg-adjusted P values were calculated separately for each comparison.

Functional analyses used Gene Ontology Biological Process (GO BP), KEGG and Reactome annotations. For over-representation analysis (ORA), proteins with P < 0.05 were separated according to the direction of their fold change and mapped to unique Entrez gene identifiers within each list. Enrichment was assessed using upper-tail hypergeometric tests, with the annotated gene universe for each collection as the background. Gene sets containing 10–500 genes were considered; sets with non-zero overlap with the input list were tested. P values were adjusted using the Benjamini– Hochberg method separately for each comparison, direction and annotation collection.

Gene set enrichment analysis (GSEA) used all mapped protein groups with valid differential statistics, ranked by −log10(P) multiplied by the sign of the log2 fold change. Ranking scores were averaged where multiple protein groups mapped to the same Entrez identifier. Analyses used fgseaMultilevel with gene-set sizes of 10–500, 10,000 initial simple permutations and eps = 10⁻¹⁰. Pathways with BH-adjusted P < 0.05 were considered significant. Selected biologically relevant terms were displayed, with complete enrichment results provided in the supplementary data.

EV proteomic data were used for descriptive characterisation of proteins detected in the isolated preparations. Selected EV-associated and macrophage-associated proteins were visualised using observed log2 LFQ intensities, and overlap between detected protein sets was summarised.

### RNA extraction from EVs and parent cells

Total RNA was extracted from macrophage cell pellets using the RNeasy Mini Kit (Qiagen) in RLT buffer, or from EV preparations using the Total Exosome RNA C Protein Isolation Kit (Thermo Fisher Scientific), according to the manufacturers’ instructions. For miRNA analysis, a synthetic spike-in (cel-miR-39; Norgen Biotek) was added during extraction for normalisation. RNA concentration and quality were assessed using a NanoDrop ND-1000 spectrophotometer (Thermo Fisher Scientific).

For gene-expression analysis, 500 ng RNA was reverse-transcribed using the QuantiTect Reverse Transcription Kit (Qiagen). miRNAs were reverse-transcribed using TaqMan Small RNA Assays (Applied Biosystems). Real-time qPCR was performed on a LightCycler 480 II (Roche) using Qiagen QuantiTect primers or TaqMan assays (Thermo Fisher Scientific) in a final volume of 10 µl. Reactions were performed in duplicate. Gene expression was normalised to β2-microglobulin or *Ppia* and miRNA expression to cel-miR-39 using the 2^−ΔΔCt method. Primer and assay details are provided in Supplementary Table 2.

### MicroRNA sequencing and analysis

Total RNA quality was assessed using an Agilent 2100 Bioanalyzer. Libraries were prepared from 100 ng cellular RNA or 5 µl EV RNA using the QIAseq miRNA Library Kit with unique dual-index barcodes and unique molecular identifiers (UMIs). Libraries were amplified using 16 PCR cycles for cellular RNA and 22 cycles for EV RNA, purified, quantified and pooled before adapter-dimer removal by gel extraction. Sequencing was performed on an Illumina NextSeq 2000 using 75-bp single-end reads with 4% PhiX.

Reads were aligned to mouse miRNAs annotated in miRBase v22, and differential-expression analysis was performed using the QIAGEN RNA-seq Analysis Portal. AAM-EVs were compared with parent AAMs to assess relative enrichment in EVs and with BMDM-EVs to identify differences associated with alternative macrophage activation. Fold changes, nominal *P* values, false discovery rate (FDR)-adjusted *P* values and counts per million (CPM)-normalised expression were exported to R. Given the small number of biological replicates (n = 3 per group), AAM-EV versus BMDM-EV comparisons were considered exploratory, with nominal *P* < 0.05 used to nominate candidate miRNAs. miR-181b-5p (*P* = 0.052) was additionally retained as a candidate based on its concordant directional difference.

For descriptive analyses, miRNAs were considered detected at CPM > 0 and retained if detected in at least two of three AAM-EV libraries. A separate exploratory dual-enrichment set comprised miRNAs with mean abundance ≥20 CPM and greater than twofold enrichment in AAM-EVs relative to both BMDM-EVs and parent AAMs.

Experimentally validated targets of miR-149-5p, miR-451a, miR-27a-3p and miR-181b-5p were retrieved using multiMiR (v1.24.1). Murine miRNAs were mapped to their corresponding human homologues to access the more extensive human validated-target annotations. Functional enrichment of validated targets was assessed using Gene Ontology Biological Process and KEGG pathway annotations.

### Single-Nuclei Isolation and Library Preparation

For the first study, mice received human alternatively activated macrophages (hAAMs). Hepatic nuclei were isolated using a protocol adapted from Carlessi et al. Briefly, approximately 25 mg of frozen liver tissue was homogenised using a Dounce tissue grinder (Sigma-Aldrich, D8938) with 15 strokes of pestle A in 2 mL of ice-cold nuclei lysis buffer (10 mM Tris-HCl, 10 mM NaCl, 3 mM MgCl2, and 0.1% IGEPAL CA-630). An additional 2 mL of lysis buffer was added, and samples were incubated on ice for 10 min before filtration through a 40 µm cell strainer. Nuclei were pelleted at 500 × g for 5 min at 4°C, washed in PBS containing 1% BSA and 0.2 U/µL RNase inhibitor, and isolated by fluorescence-activated cell sorting (FACS) following 7-AAD staining. Approximately 100,000 nuclei were collected per sample. Single-nuclei libraries were prepared using the 10x Genomics Chromium Single Cell 3′ Gene Expression v3.1 chemistry, dual-indexed (SI-TT-H2 to H9), and cDNA was amplified using 11 PCR cycles. Sequencing was performed on an Illumina NextSeq 2000 instrument using a P3 flow cell (100 cycles).

For the second study, mice received either mouse AAMs or AAM-derived extracellular vesicles (AAM-EVs). Nuclei were isolated using a Tween-salt-Tris (TST)-based method. Approximately 45 mg of frozen liver tissue was minced in 1 mL of ice-cold TST buffer containing 146 mM NaCl, 10 mM Tris-HCl (pH 7.5), 1 mM CaCl2, 21 mM MgCl2, 0.03% Tween-20, 0.01% BSA, and RNase inhibitor (N2615, Promega). Tissue was mechanically dissociated on ice for 10 min with micro-scissors, passed through a 40 µm cell strainer, and washed with an additional 1 mL of TST buffer. Lysis was arrested by adding 3 mL of chilled salt-Tris buffer without detergent. Nuclei were pelleted at 500 × g for 5 min at 4°C, resuspended in PBS containing BSA and RNase inhibitor, and filtered through a 35 µm strainer. DAPI-positive nuclei were isolated via FACS, with approximately 100,000 nuclei collected per sample. Following quality assessment of nuclear integrity and concentration, freshly isolated nuclei were loaded onto the 10x Genomics Chromium platform. Libraries were prepared using Chromium Single Cell 3′ Gene Expression v4 chemistry, dual-indexed (SI-TT-G4 to G12), with 11 PCR cycles of cDNA amplification. Initial quality-control sequencing was performed using an Illumina iSeq 100, followed by high-depth production sequencing on a NextSeq 2000 instrument using a P4 flow cell (100 cycles). Library quality in both studies was validated using an Agilent Bioanalyzer and Qubit fluorometry.

### Preprocessing and Quality Control of snRNA-seq Data

Raw sequencing data were processed using Cell Ranger (10x Genomics; v6.1.2 for study one, v7.1.0 for study two) and aligned to the 10x Genomics mm10-2020-A reference genome. Alignment and barcode counting included intronic reads to account for pre-mRNA in nascent nuclear transcripts. For study one, sequencing yielded 2,000–5,100 nuclei and 229–394 million reads per sample (mean depth: 52,159–195,293 reads/nucleus; median detection: 521–3,257 genes/nucleus; saturation up to 91%; confident transcriptome mapping: 64.7%–70.0%). For study two, sequencing yielded 6,731–36,876 nuclei and 139–239 million reads per sample (mean depth: 6,731–27,619 reads/nucleus; median detection: 1,287–3,012 genes/nucleus; saturation: 15.9%– 32.0%; confident transcriptome mapping: 73.6%–85.6%).

Filtered count matrices from Cell Ranger were imported into R for downstream processing via Seurat. Initial filtering retained features detected in ≥ 3 cells and cells with ≥ 200 detected features. Ambient mRNA contamination was estimated and computationally removed using SoupX (v1.6.2). Decontaminated count matrices were then re-filtered using the same baseline parameters to remove lowly expressed features, and cells exhibiting > 5% mitochondrial reads were excluded. Corrected samples were merged, normalized, scaled, and subjected to batch correction using Harmony (v1.2.3) with 30 iterations. Doublets were predicted and eliminated using scDblFinder (v1.20.2), and cell clusters displaying high predicted doublet scores were discarded. An initial estimate of cell types was computed using SingleR (v2.8.0) using references from previously characterised liver snRNA-seq data. Final cell type identities were manually assigned to clusters based on SingleR predictions and canonical lineage markers, and clusters were cleaned of contaminating nuclei using a supervised verification approach.

### Dimensionality Reduction, Clustering, and Differential Expression

Dimensionality reduction was performed using Principal Component Analysis (PCA) followed by Uniform Manifold Approximation and Projection (UMAP), with cell populations identified via graph-based Louvain clustering. To assess treatment-associated differential expression within annotated populations, single-cell RNA counts were corrected using SoupX, aggregated by individual mouse to generate animal-level pseudobulk profiles, and filtered for low expression. Differential expression was evaluated using edgeR with TMM normalisation and quasi-likelihood testing, with limma-voom serving as a complementary validation framework. In the three-condition study, we also used MAST in Seurat to screen for treatment-associated differences in log-normalised RNA expression within annotated myeloid populations, including a targeted analysis of cycling genes within each Kupffer cell state. Nuclei were pooled across animals within each condition, and AAM and AAM-EV were each compared with PBS. Models included the number of genes detected per nucleus as a covariate. Given the limited biological replicates and inter-animal variability, nominal *P* < 0.05 was used to identify candidates for downstream pathway analysis. Functional over-representation analysis was conducted using clusterProfiler across Gene Ontology Biological Process, KEGG, and Reactome annotations, with pathway-level P values adjusted using the Benjamini–Hochberg method.

### Statistics

Statistical analyses were performed using GraphPad Prism 9.0 (GraphPad Software, La Jolla, CA) or R, as appropriate. Data were first assessed for normality and equal variance to determine the appropriate statistical test. For normally distributed data with equal variances, one-way analysis of variance (ANOVA) was used, with Brown–Forsythe and Welch correction applied for multiple comparisons when comparing across time points within the same organ. For non-normally distributed data, the Kruskal–Wallis test was used, followed by Dunn’s post hoc test for group-wise comparisons. For the phagocytosis time course (Fig. 2D), the fraction of pHrodo-positive cells was measured repeatedly in the same wells and analysed by two-way repeated-measures ANOVA, with treatment as the between-well factor and time as the within-well factor. A Geisser–Greenhouse correction was applied, followed by Dunnett’s comparisons of each treatment with PBS at each time point.

### Study approval

All animal experiments were undertaken in accordance with criteria outlined in a license granted under the Animals (Scientific Procedures) Act 1986 and approved by the University of Edinburgh Animal Ethics Committee. This study was conducted in compliance with all relevant ethical regulations, including the principles outlined in the Declaration of Helsinki. The use of human material was granted by the South East Scotland Research Ethics Committee 02, and the use of buffy coats was covered by the Scottish National Blood Transfusion Service (SNBTS). Buffy coats from informed consenting healthy volunteers were obtained in collaboration with SNBTS Blood Donor Centre, Edinburgh, United Kingdom, under SNBTS Sample Governance 16-09.

