## Supplementary data for "Extracellular vesicles from alternatively activated macrophages remodel the injured hepatic niche and promote repair"

Fig S1

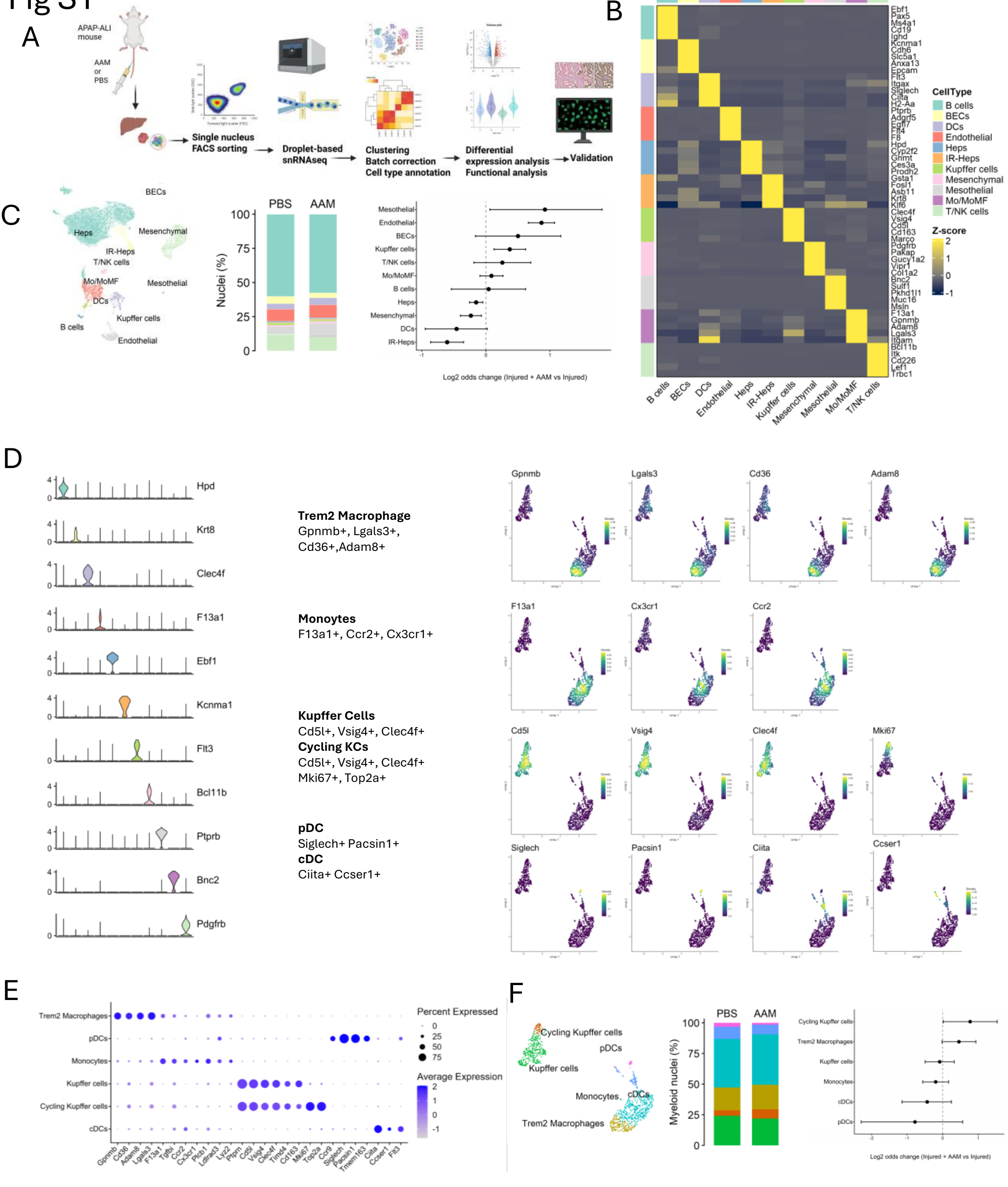
A
APAP-ALI mouse
AAM or PBS
Single nucleus FACS sorting
Droplet-based snRNAseq
Clustering
Batch correction
Cell type annotation
Differential expression analysis
Functional analysis
Validation
B
CellType
B cells
BECs
DCs
Endothelial
Heps
IR-Heps
Kupffer cells
Mesenchymal
Mesothelial
Mo/MoMF
T/NK cells
Z-score
C
PBS
AAM
Nuclei (%)
Log2 odds change (Injured + AAM vs Injured)
D
Trem2 Macrophage
Gpnmb+, Lgals3+, Cd36+,Adam8+
Monoytes
F13a1+, Ccr2+, Cx3cr1+
Kupffer Cells
Cd5l+, Vsig4+, Clec4f+
Cycling KCs
Cd5l+, Vsig4+, Clec4f+ Mki67+, Top2a+
pDC
Siglech+ Pacsin1+
cDC
Ciita+ Ccser1+
E
Percent Expressed
Average Expression
Trem2 Macrophages
pDCs
Monocytes
Kupffer cells
Cycling Kupffer cells
cDCs
F
Cycling Kupffer cells
pDCs
Kupffer cells
Monocytes
cDCs
Trem2 Macrophages
Myeloid nuclei (%)
Log2 odds change (Injured + AAM vs Injured)

Fig S2

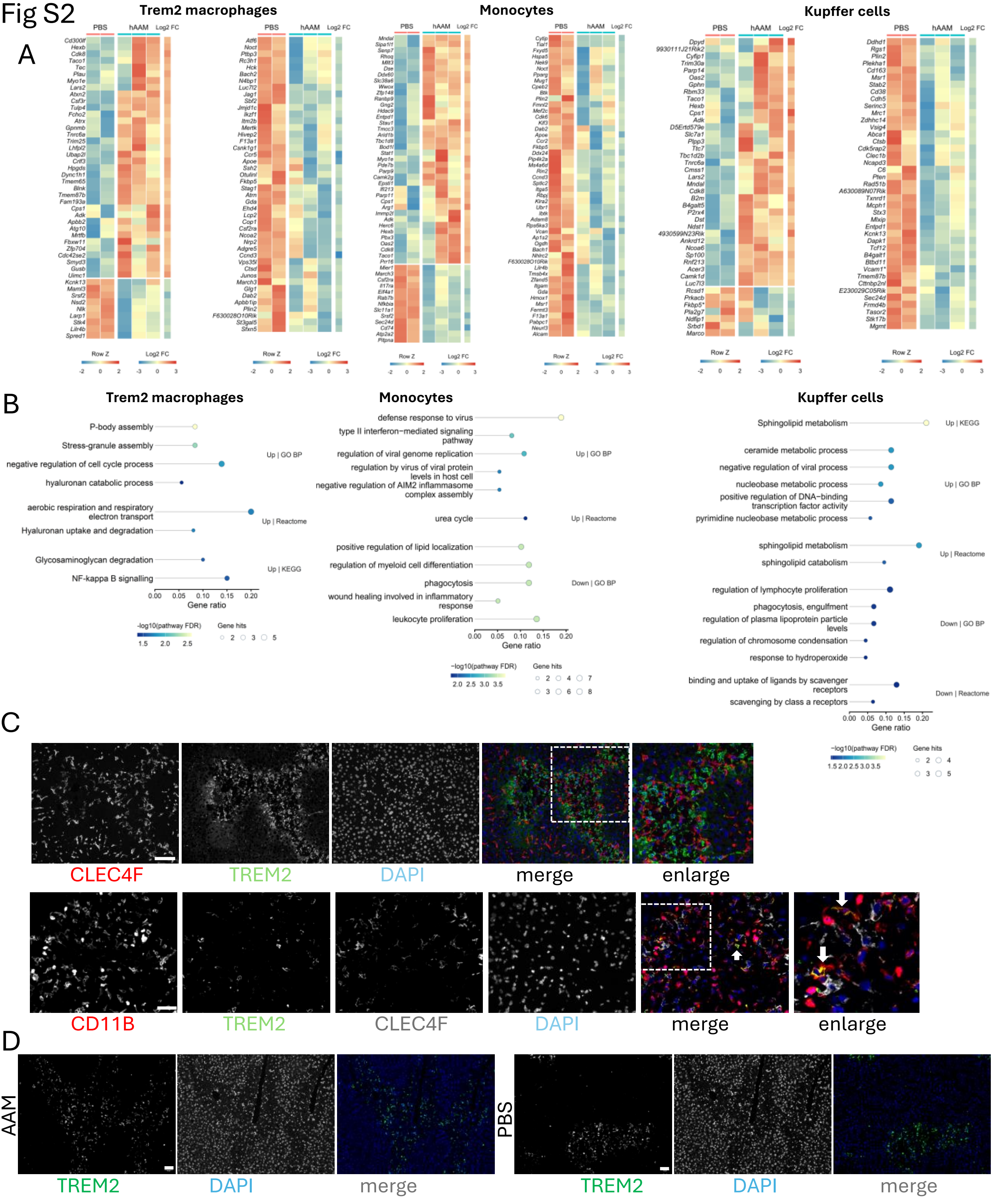
A
Trem2 macrophages
Monocytes
Kupffer cells
PBS
hAAM
Log2 FC
Row Z
B
Trem2 macrophages
P-body assembly
Stress-granule assembly
negative regulation of cell cycle process
hyaluronan catabolic process
Up | GO BP
aerobic respiration and respiratory electron transport
Hyaluronan uptake and degradation
Up | Reactome
Glycosaminoglycan degradation
NF-kappa B signalling
Up | KEGG
Gene ratio
-log10(pathway FDR)
Gene hits
Monocytes
defense response to virus
type II interferon−mediated signaling pathway
regulation of viral genome replication
regulation by virus of viral protein levels in host cell
negative regulation of AIM2 inflammasome complex assembly
Up | GO BP
urea cycle
Up | Reactome
positive regulation of lipid localization
regulation of myeloid cell differentiation
phagocytosis
wound healing involved in inflammatory response
leukocyte proliferation
Down | GO BP
Kupffer cells
Sphingolipid metabolism
Up | KEGG
ceramide metabolic process
negative regulation of viral process
nucleobase metabolic process
positive regulation of DNA−binding transcription factor activity
pyrimidine nucleobase metabolic process
Up | GO BP
sphingolipid metabolism
sphingolipid catabolism
Up | Reactome
regulation of lymphocyte proliferation
phagocytosis, engulfment
regulation of plasma lipoprotein particle levels
regulation of chromosome condensation
response to hydroperoxide
Down | GO BP
binding and uptake of ligands by scavenger receptors
scavenging by class a receptors
Down | Reactome
C
CLEC4F
TREM2
DAPI
merge
enlarge
CD11B
TREM2
CLEC4F
DAPI
merge
enlarge
D
AAM
TREM2
DAPI
merge
PBS
TREM2
DAPI
merge

Fig S3

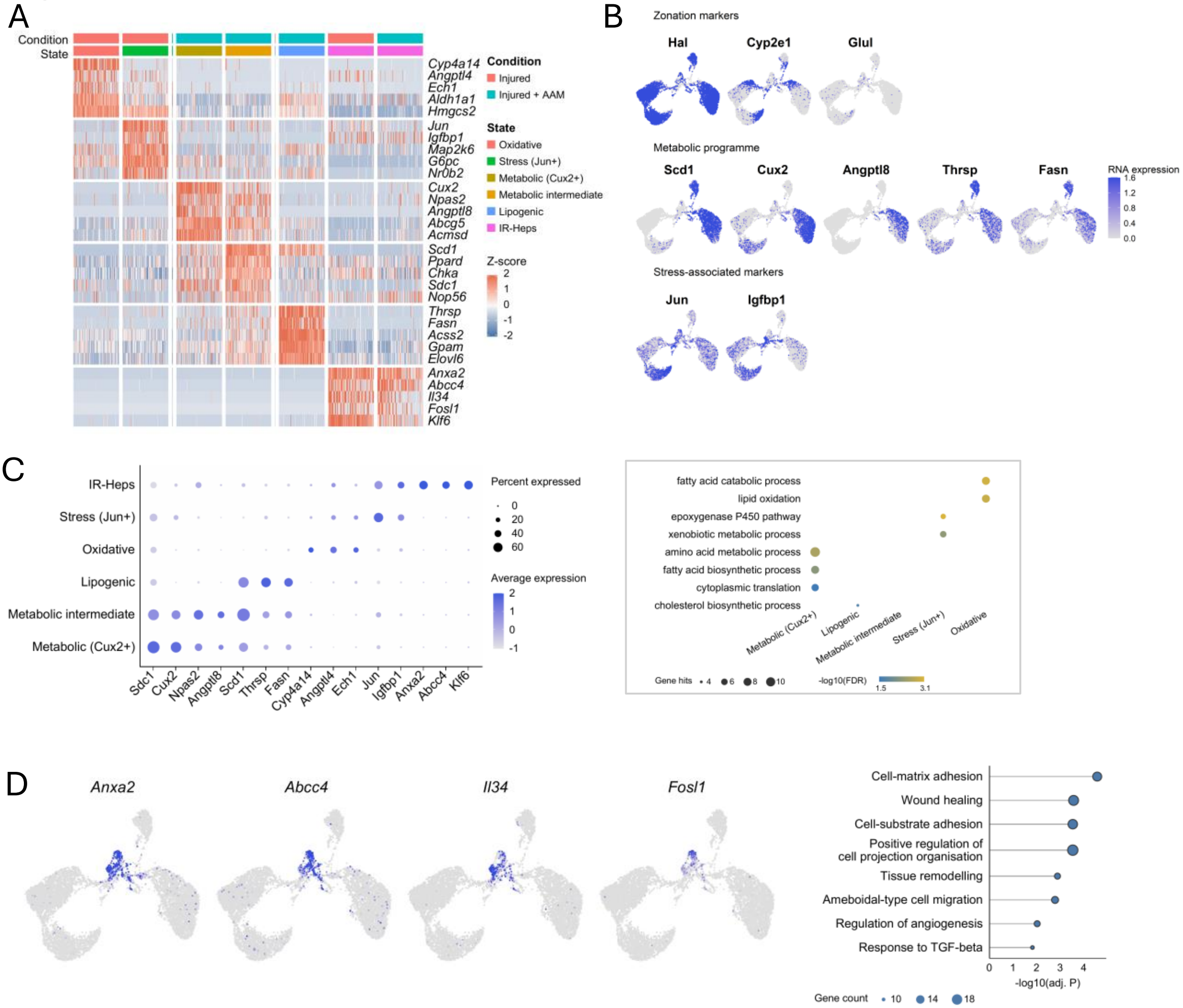
A
Condition
State
Cyp4a14
Angptl4
Ech1
Aldh1a1
Hmgcs2
Jun
Igfbp1
Map2k6
G6pc
Nr0b2
Cux2
Npas2
Angptl8
Abcg5
Acmsd
Scd1
Ppard
Chka
Sdc1
Nop56
Thrsp
Fasn
Acss2
Gpam
Elovl6
Anxa2
Abcc4
Il34
Fosl1
Klf6
Condition
Injured
Injured + AAM
State
Oxidative
Stress (Jun+)
Metabolic (Cux2+)
Metabolic intermediate
Lipogenic
IR-Heps
Z-score
2
1
0
-1
-2
B
Zonation markers
Hal
Cyp2e1
Glul
Metabolic programme
Scd1
Cux2
Angptl8
Thrsp
Fasn
RNA expression
1.6
1.2
0.8
0.4
0.0
Stress-associated markers
Jun
Igfbp1
C
IR-Heps
Stress (Jun+)
Oxidative
Lipogenic
Metabolic intermediate
Metabolic (Cux2+)
Sdc1
Cux2
Npas2
Angptl8
Scd1
Thrsp
Fasn
Cyp4a14
Angptl4
Ech1
Jun
Igfbp1
Anxa2
Abcc4
Klf6
Percent expressed
0
20
40
60
Average expression
2
1
0
-1
fatty acid catabolic process
lipid oxidation
epoxygenase P450 pathway
xenobiotic metabolic process
amino acid metabolic process
fatty acid biosynthetic process
cytoplasmic translation
cholesterol biosynthetic process
Metabolic (Cux2+)
Lipogenic
Metabolic intermediate
Stress (Jun+)
Oxidative
Gene hits
4
6
8
10
-log10(FDR)
1.5
3.1
D
Anxa2
Abcc4
Il34
Fosl1
Cell-matrix adhesion
Wound healing
Cell-substrate adhesion
Positive regulation of cell projection organisation
Tissue remodelling
Ameboidal-type cell migration
Regulation of angiogenesis
Response to TGF-beta
0
1
2
3
4
-log10(adj. P)
Gene count
10
14
18

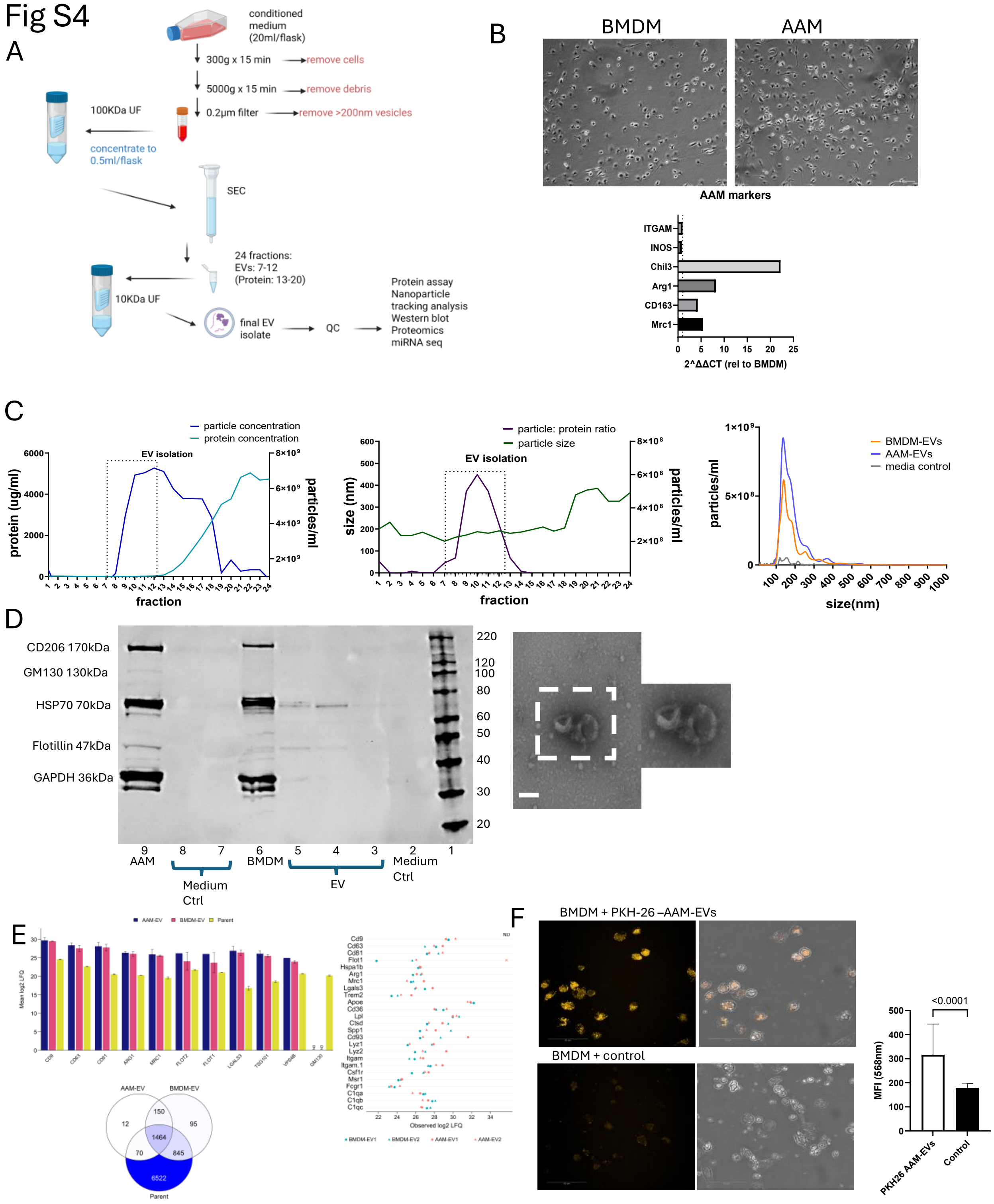

Fig S4
A
conditioned medium (20ml/flask)
300g x 15 min
remove cells
5000g x 15 min
remove debris
0.2µm filter
remove >200nm vesicles
100KDa UF
concentrate to 0.5ml/flask
SEC
24 fractions: EVs: 7-12 (Protein: 13-20)
10KDa UF
final EV isolate
QC
Protein assay
Nanoparticle tracking analysis
Western blot
Proteomics
miRNA seq
B
BMDM
AAM
AAM markers
ITGAM
INOS
Chil3
Arg1
CD163
Mrc1
2^ΔΔCT (rel to BMDM)
C
particle concentration
protein concentration
EV isolation
protein (ug/ml)
particles/ml
fraction
particle: protein ratio
particle size
size (nm)
BMDM-EVs
AAM-EVs
media control
size(nm)
D
CD206 170kDa
GM130 130kDa
HSP70 70kDa
Flotillin 47kDa
GAPDH 36kDa
AAM
BMDM
Medium Ctrl
EV
E
AAM-EV
BMDM-EV
Parent
Mean log2 LFQ
Observed log2 LFQ
F
BMDM + PKH-26 –AAM-EVs
BMDM + control
<0.0001
MFI (568nm)
PKH26 AAM-EVs
Control

Fig S5

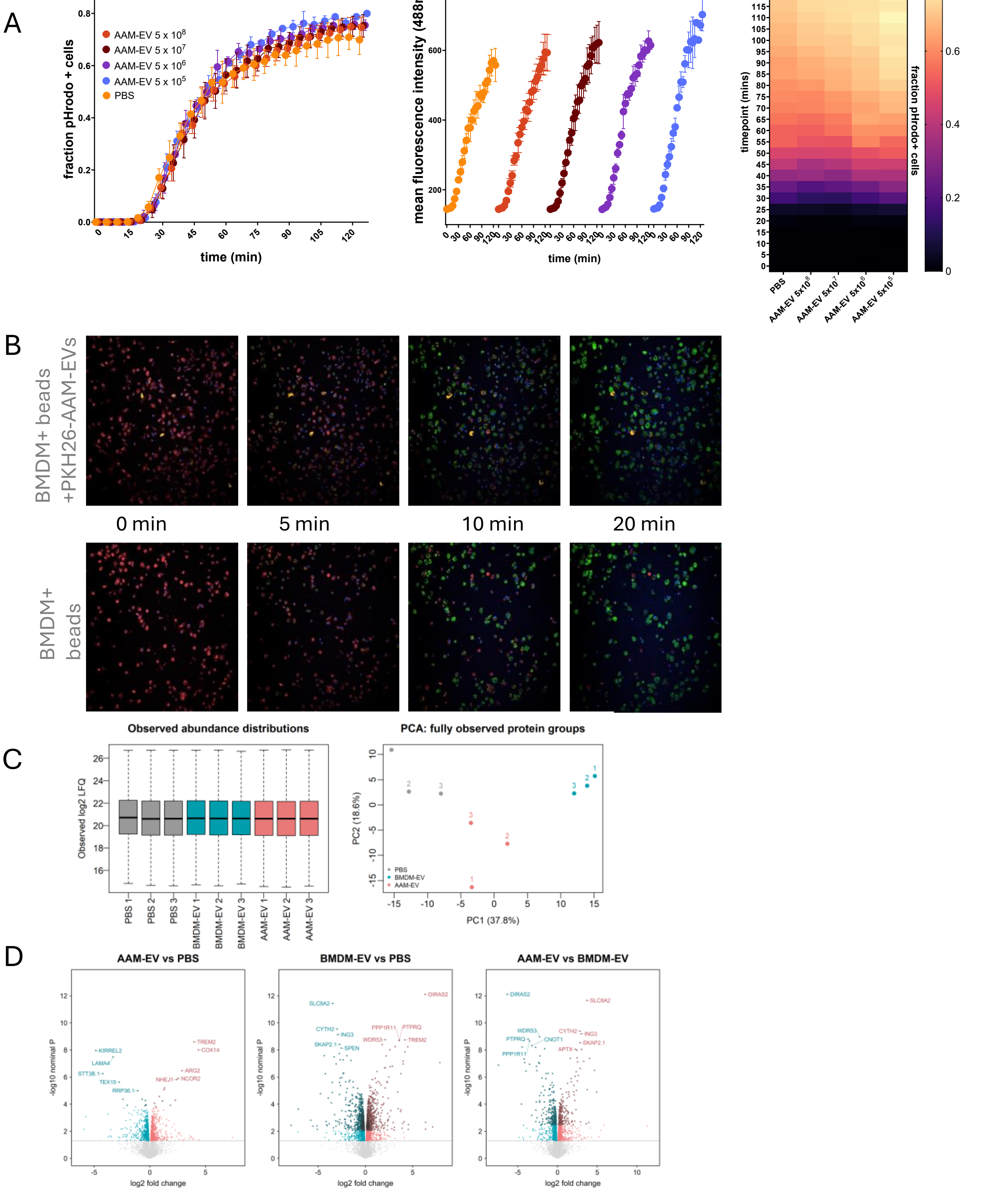
A
fraction pHrodo + cells
AAM-EV 5 x 10^8
AAM-EV 5 x 10^7
AAM-EV 5 x 10^6
AAM-EV 5 x 10^5
PBS
time (min)
mean fluorescence intensity (488nm)
time (min)
timepoint (mins)
fraction pHrodo+ cells
PBS
AAM-EV 5x10^8
AAM-EV 5x10^7
AAM-EV 5x10^6
AAM-EV 5x10^5
B
BMDM+ beads +PKH26-AAM-EVs
0 min
5 min
10 min
20 min
BMDM+ beads
C
Observed abundance distributions
Observed log2 LFQ
PBS 1
PBS 2
PBS 3
BMDM-EV 1
BMDM-EV 2
BMDM-EV 3
AAM-EV 1
AAM-EV 2
AAM-EV 3
PCA: fully observed protein groups
PC1 (37.8%)
PC2 (18.6%)
PBS
BMDM-EV
AAM-EV
D
AAM-EV vs PBS
BMDM-EV vs PBS
AAM-EV vs BMDM-EV
-log10 nominal P
log2 fold change
KIRREL2
LAMA4
STT3B.1
TEX15
RRP36.1
TREM2
COX14
ARG2
NCOR2
NHEJ1
DIRAS2
SLC8A2
CYTH2
ING3
SKAP2.1
SPEN
PPP1R11
PTPRQ
WDR53
CNOT1
APTX

Fig S6

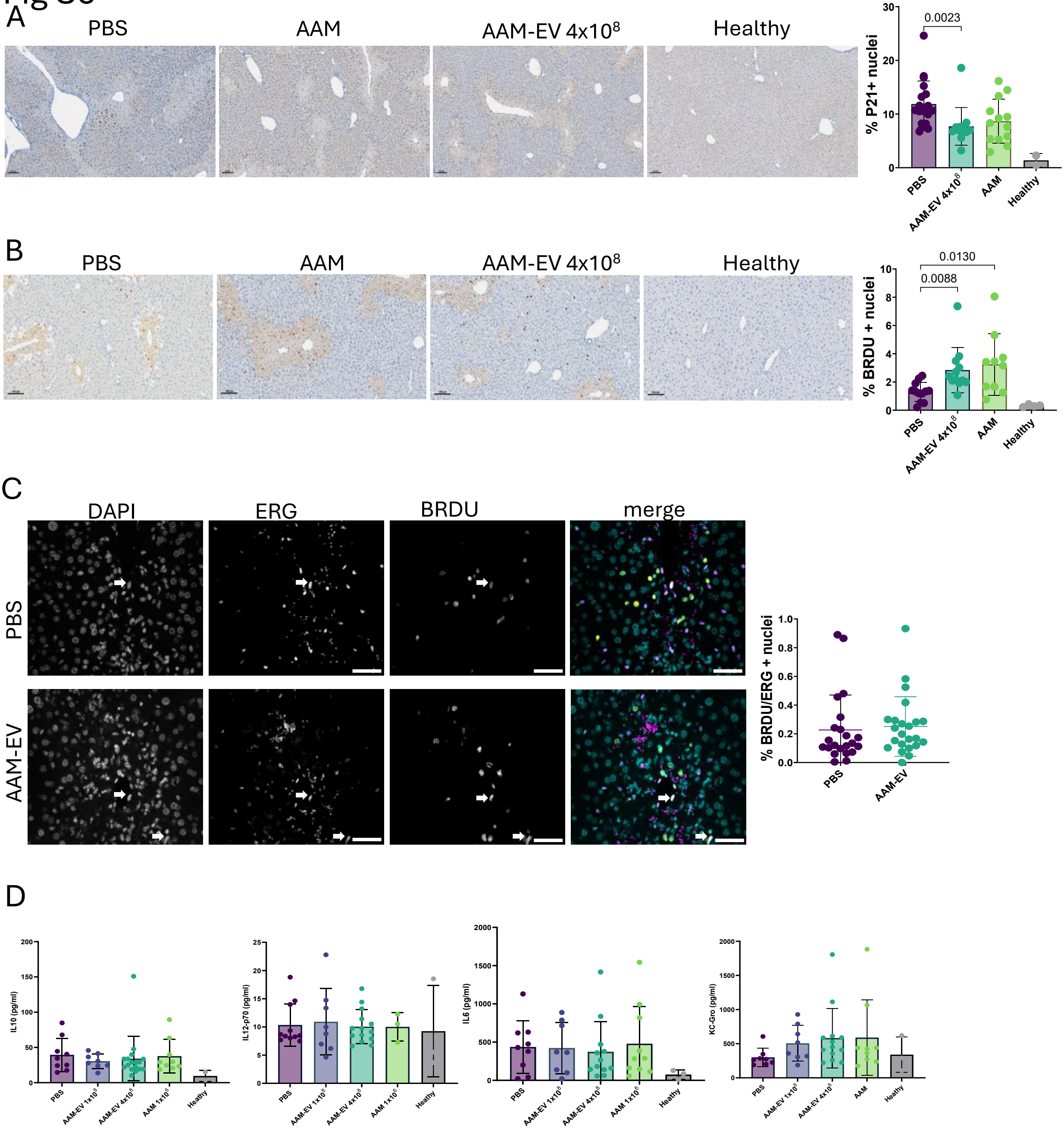
A
PBS
AAM
AAM-EV 4x10⁸
Healthy
% P21+ nuclei
0.0023
PBS
AAM-EV 4x10⁸
AAM
Healthy
B
PBS
AAM
AAM-EV 4x10⁸
Healthy
% BRDU + nuclei
0.0130
0.0088
PBS
AAM-EV 4x10⁸
AAM
Healthy
C
DAPI
ERG
BRDU
merge
PBS
AAM-EV
% BRDU/ERG + nuclei
PBS
AAM-EV
D
IL10 (pg/ml)
IL12-p70 (pg/ml)
IL6 (pg/ml)
KC-Gro (pg/ml)
PBS
AAM-EV 1x10⁸
AAM-EV 4x10⁸
AAM 1x10⁶
AAM
Heathy

### Fig S7

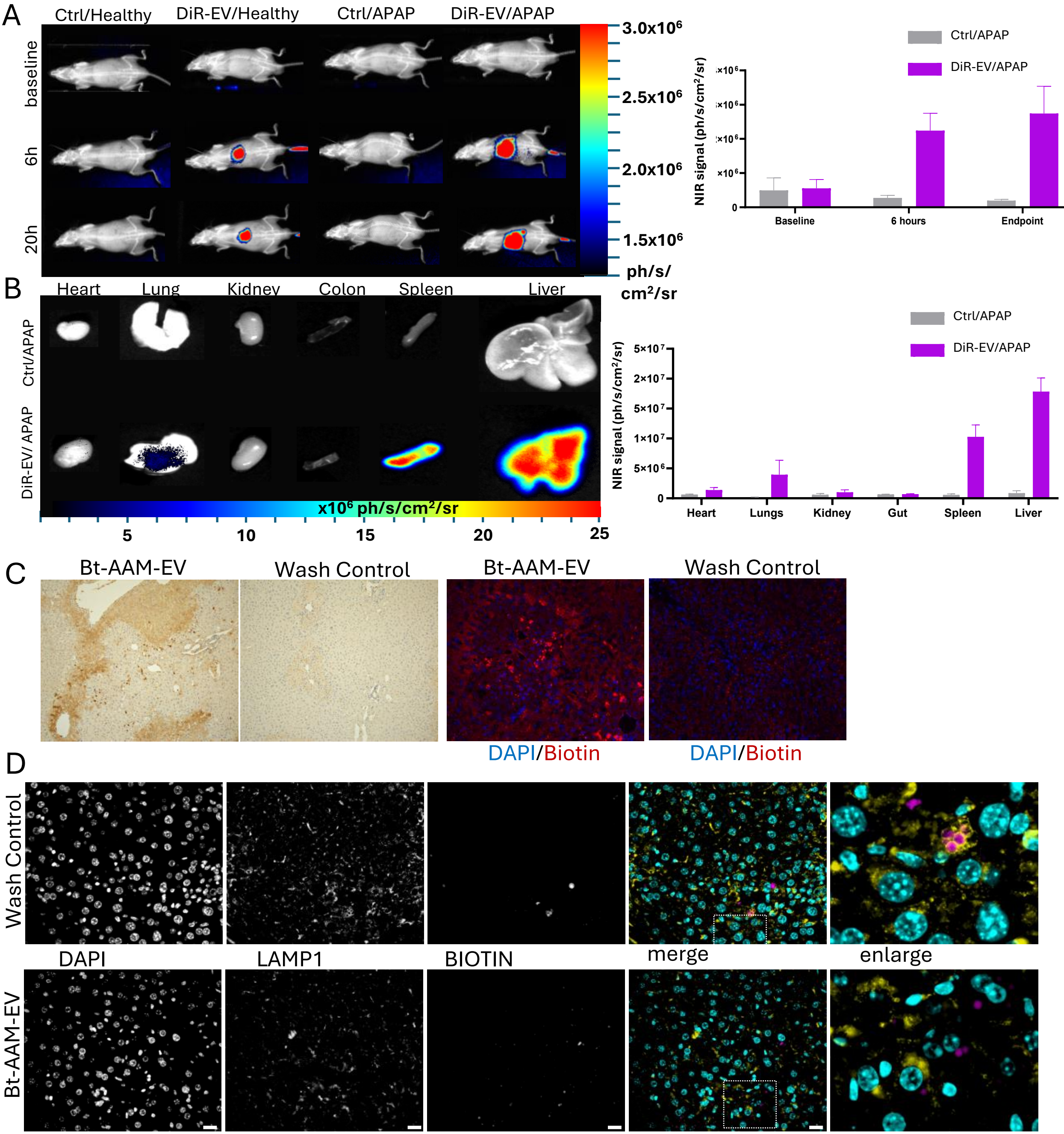

A
Ctrl/Healthy
DiR-EV/Healthy
Ctrl/APAP
DiR-EV/APAP
baseline
6h
20h
3.0x10^6
2.5x10^6
2.0x10^6
1.5x10^6
ph/s/cm²/sr
Ctrl/APAP
DiR-EV/APAP
NIR signal (ph/s/cm²/sr)
Baseline
6 hours
Endpoint
B
Heart
Lung
Kidney
Colon
Spleen
Liver
Ctrl/APAP
DiR-EV/ APAP
x10^6 ph/s/cm²/sr
5
10
15
20
25
5×10^7
2×10^7
5×10^7
1×10^7
5×10^6
0
Heart
Lungs
Kidney
Gut
Spleen
Liver
C
Bt-AAM-EV
Wash Control
DAPI/Biotin
D
Wash Control
Bt-AAM-EV
DAPI
LAMP1
BIOTIN
merge
enlarge

### Fig S8

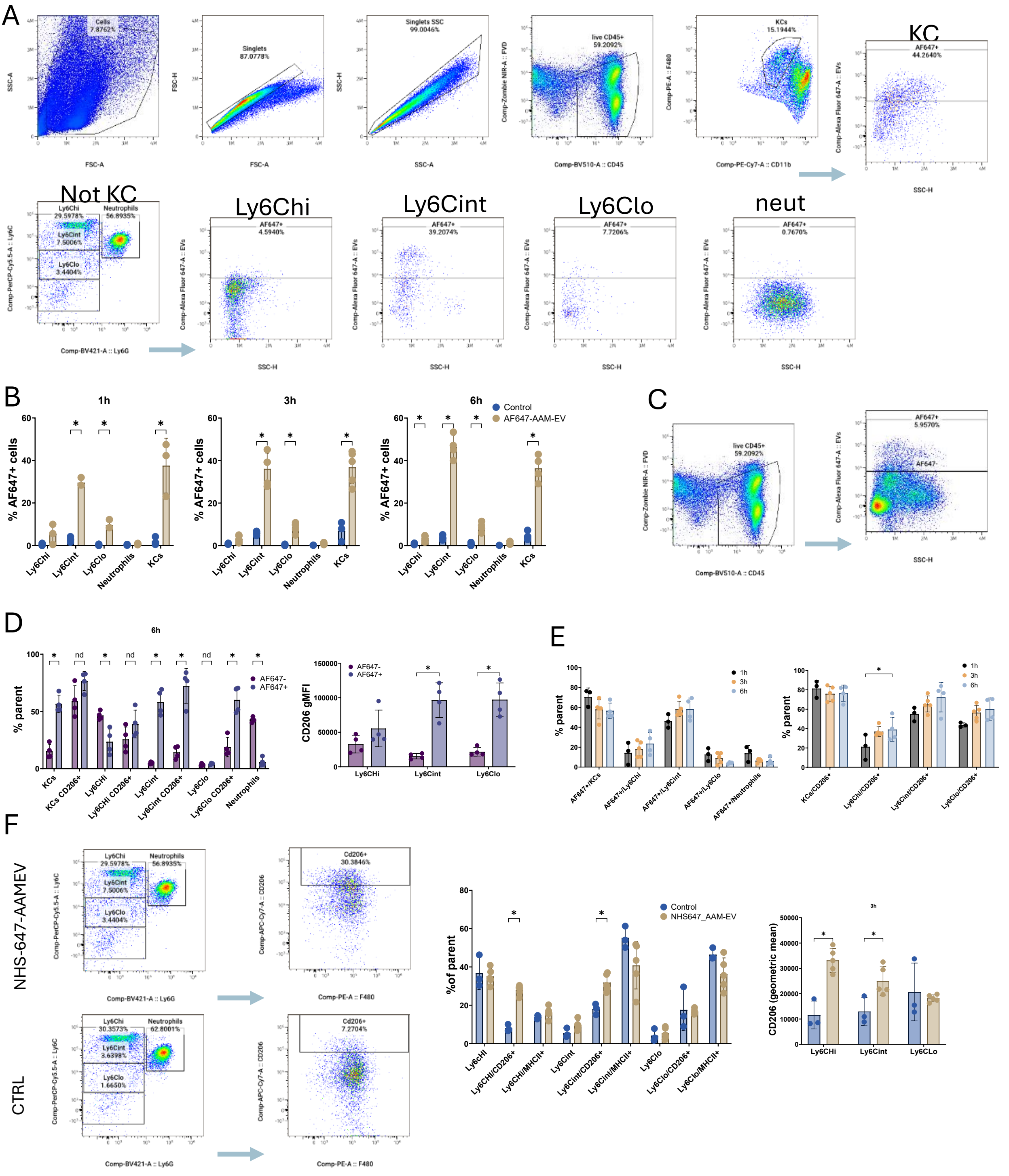
A
KC
Not KC
Ly6Chi
Ly6Cint
Ly6Clo
neut
B
1h
3h
6h
Control
AF647-AAM-EV
% AF647+ cells
C
D
6h
% parent
AF647-
AF647+
CD206 gMFI
E
1h
3h
6h
F
NHS-647-AAMEV
CTRL
Control
NHS647_AAM-EV
%of parent
CD206 (geometric mean)

### Fig S9

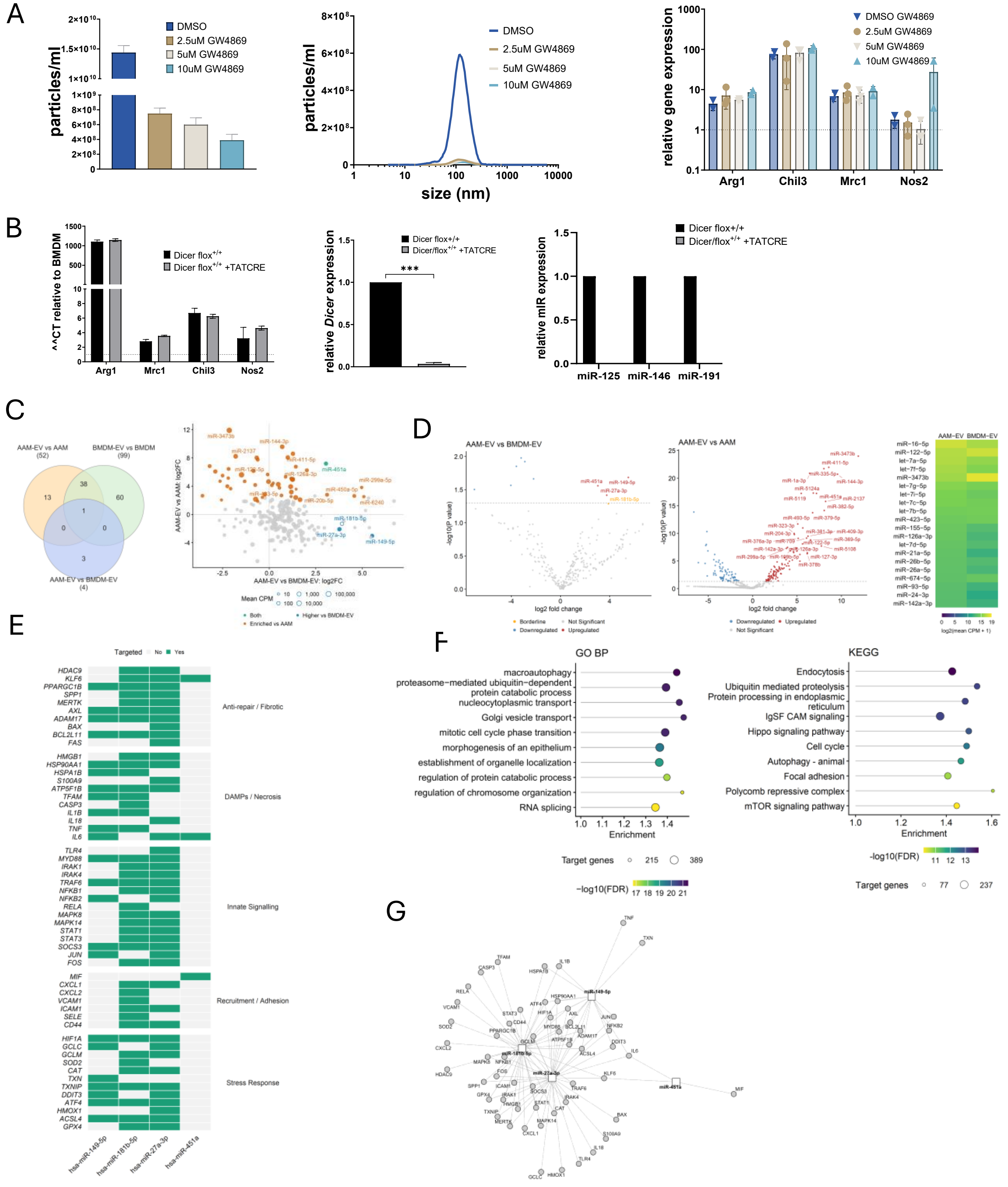

A
particles/ml
DMSO
2.5uM GW4869
5uM GW4869
10uM GW4869
size (nm)
relative gene expression
DMSO GW4869
Arg1
Chil3
Mrc1
Nos2
B
^^CT relative to BMDM
Dicer flox+/+
Dicer flox+/+ +TATCRE
relative Dicer expression
Dicer/flox+/+ +TATCRE
relative mIR expression
miR-125
miR-146
miR-191
C
AAM-EV vs AAM (52)
BMDM-EV vs BMDM (99)
AAM-EV vs BMDM-EV (4)
AAM-EV vs BMDM-EV: log2FC
AAM-EV vs AAM: log2FC
D
AAM-EV vs BMDM-EV
AAM-EV vs AAM
log2 fold change
-log10(P value)
AAM-EV
BMDM-EV
log2(mean CPM + 1)
E
Targeted
Anti-repair / Fibrotic
DAMPs / Necrosis
Innate Signalling
Recruitment / Adhesion
Stress Response
hsa-miR-149-5p
hsa-miR-181b-5p
hsa-miR-27a-3p
hsa-miR-451a
F
GO BP
macroautophagy
proteasome−mediated ubiquitin−dependent protein catabolic process
nucleocytoplasmic transport
Golgi vesicle transport
mitotic cell cycle phase transition
morphogenesis of an epithelium
establishment of organelle localization
regulation of protein catabolic process
regulation of chromosome organization
RNA splicing
Enrichment
Target genes
-log10(FDR)
KEGG
Endocytosis
Ubiquitin mediated proteolysis
Protein processing in endoplasmic reticulum
IgSF CAM signaling
Hippo signaling pathway
Cell cycle
Autophagy - animal
Focal adhesion
Polycomb repressive complex
mTOR signaling pathway
G

Fig S10

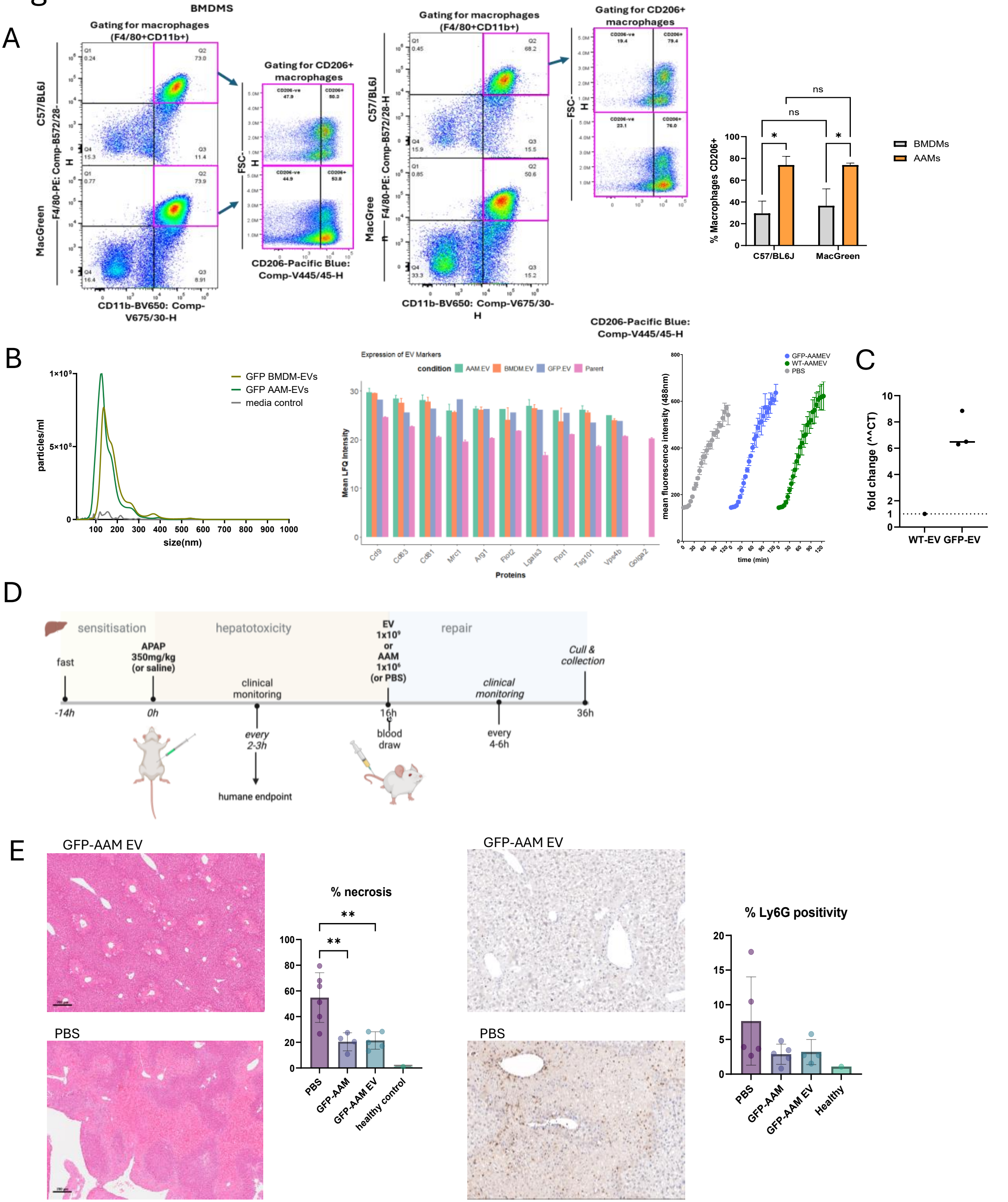
A
BMDMS
Gating for macrophages (F4/80+CD11b+)
Gating for CD206+ macrophages
C57/BL6J
MacGreen
F4/80-PE: Comp-B572/28-H
CD11b-BV650: Comp-V675/30-H
CD206-Pacific Blue: Comp-V445/45-H
FSC-H
AAMs
% Macrophages CD206+
BMDMs
AAMs
ns
*
B
1×10⁹
5×10⁸
particles/ml
size(nm)
GFP BMDM-EVs
GFP AAM-EVs
media control
Expression of EV Markers
condition
AAM.EV
BMDM.EV
GFP.EV
Parent
Mean LFQ Intensity
Proteins
Cd9
Cd63
Cd81
Mrc1
Arg1
Flot2
Lgals3
Flot1
Tsg101
Vps4b
Golga2
GFP-AAMEV
WT-AAMEV
PBS
mean fluorescence intensity (488nm)
time (min)
C
fold change (^^CT)
WT-EV
GFP-EV
D
sensitisation
hepatotoxicity
repair
fast
APAP 350mg/kg (or saline)
clinical monitoring
EV 1x10⁹ or AAM 1x10⁶ (or PBS)
Cull & collection
-14h
0h
16h
36h
every 2-3h
humane endpoint
blood draw
every 4-6h
E
GFP-AAM EV
PBS
% necrosis
**
PBS
GFP-AAM
GFP-AAM EV
healthy control
% Ly6G positivity
Healthy

Fig S11

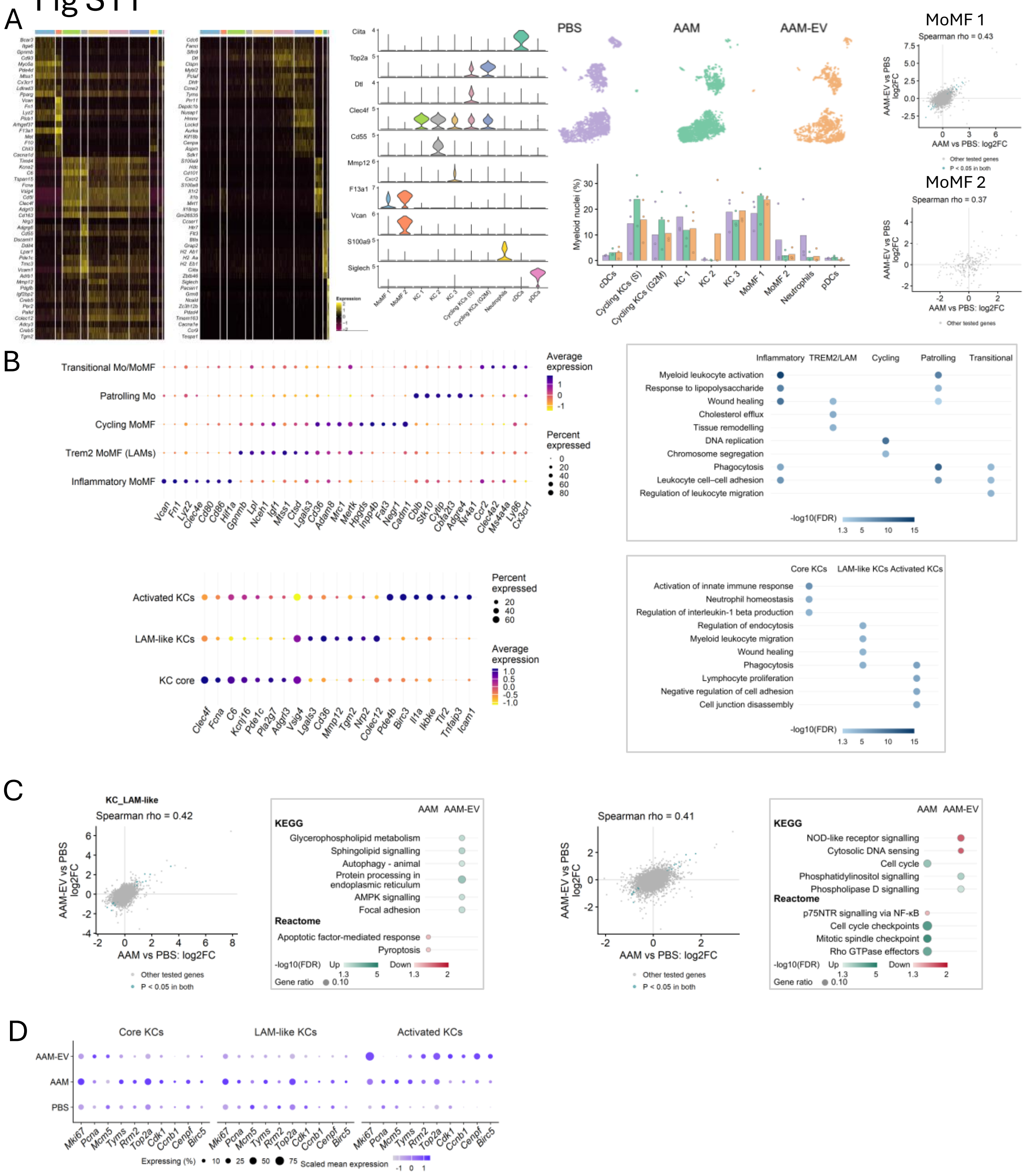
A
PBS
AAM
AAM-EV
MoMF 1
Spearman rho = 0.43
AAM-EV vs PBS log2FC
AAM vs PBS: log2FC
Other tested genes
P < 0.05 in both
MoMF 2
Spearman rho = 0.37
Myeloid nuclei (%)
cDCs
Cycling KCs (S)
Cycling KCs (G2M)
KC 1
KC 2
KC 3
MoMF 1
MoMF 2
Neutrophils
pDCs
B
Transitional Mo/MoMF
Patrolling Mo
Cycling MoMF
Trem2 MoMF (LAMs)
Inflammatory MoMF
Average expression
Percent expressed
Inflammatory
TREM2/LAM
Cycling
Patrolling
Transitional
Myeloid leukocyte activation
Response to lipopolysaccharide
Wound healing
Cholesterol efflux
Tissue remodelling
DNA replication
Chromosome segregation
Phagocytosis
Leukocyte cell–cell adhesion
Regulation of leukocyte migration
-log10(FDR)
Activated KCs
LAM-like KCs
KC core
Core KCs
Activation of innate immune response
Neutrophil homeostasis
Regulation of interleukin-1 beta production
Regulation of endocytosis
Myeloid leukocyte migration
Lymphocyte proliferation
Negative regulation of cell adhesion
Cell junction disassembly
C
KC_LAM-like
Spearman rho = 0.42
Spearman rho = 0.41
AAM
AAM-EV
KEGG
Glycerophospholipid metabolism
Sphingolipid signalling
Autophagy - animal
Protein processing in endoplasmic reticulum
AMPK signalling
Focal adhesion
Reactome
Apoptotic factor-mediated response
Pyroptosis
-log10(FDR) Up Down
Gene ratio 0.10
NOD-like receptor signalling
Cytosolic DNA sensing
Cell cycle
Phosphatidylinositol signalling
Phospholipase D signalling
p75NTR signalling via NF-κB
Cell cycle checkpoints
Mitotic spindle checkpoint
Rho GTPase effectors
D
Core KCs
LAM-like KCs
Activated KCs
AAM-EV
AAM
PBS
Mki67
Pcna
Mcm5
Tyms
Rrm2
Top2a
Cdk1
Ccnb1
Cenpf
Birc5
Expressing (%)
Scaled mean expression

### Fig S12

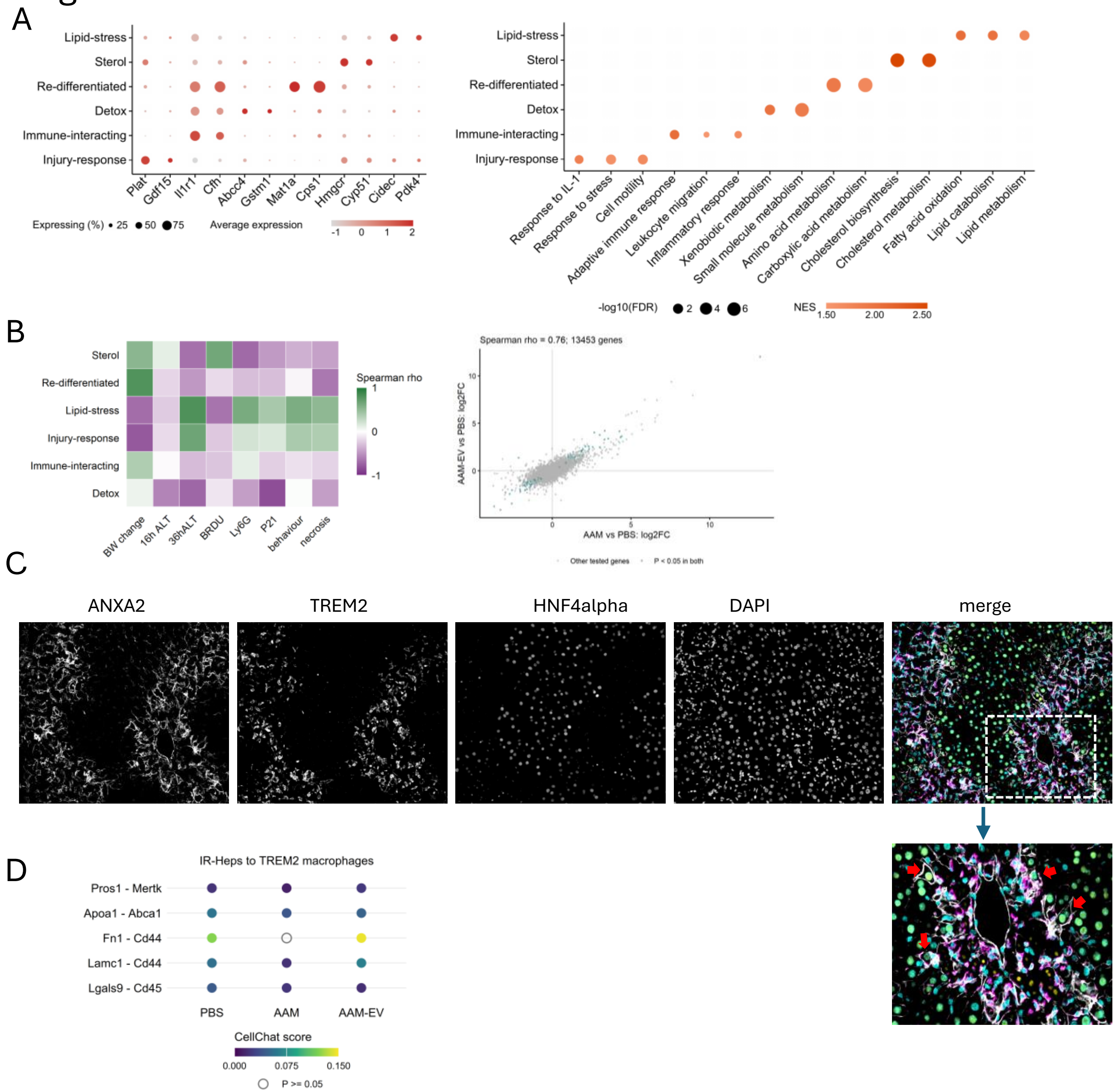

A
Lipid-stress
Sterol
Re-differentiated
Detox
Immune-interacting
Injury-response
Plat
Gdf15
Il1r1
Cfh
Abcc4
Gstm1
Mat1a
Cps1
Hmgcr
Cyp51
Cidec
Pdk4
Expressing (%) 25 50 75
Average expression
-1 0 1 2
Response to IL-1
Response to stress
Cell motility
Adaptive immune response
Leukocyte migration
Inflammatory response
Xenobiotic metabolism
Small molecule metabolism
Amino acid metabolism
Carboxylic acid metabolism
Cholesterol biosynthesis
Cholesterol metabolism
Fatty acid oxidation
Lipid catabolism
Lipid metabolism
-log10(FDR) 2 4 6
NES 1.50 2.00 2.50
B
Sterol
Re-differentiated
Lipid-stress
Injury-response
Immune-interacting
Detox
BW change
16h ALT
36hALT
BRDU
Ly6G
P21
behaviour
necrosis
Spearman rho
1
0
-1
Spearman rho = 0.76; 13453 genes
AAM-EV vs PBS: log2FC
AAM vs PBS: log2FC
Other tested genes
P < 0.05 in both
C
ANXA2
TREM2
HNF4alpha
DAPI
merge
D
IR-Heps to TREM2 macrophages
Pros1 - Mertk
Apoa1 - Abca1
Fn1 - Cd44
Lamc1 - Cd44
Lgals9 - Cd45
PBS
AAM
AAM-EV
CellChat score
0.000
0.075
0.150
P >= 0.05

**Figure S1. A single nuclei atlas of APAP-induced acute liver injury following hAAM therapy. Characterization of liver cell type in injured (n=6) or healthy mice(n=2) receiving hAAMs or PBS.**

A) Schematic illustrating workflow for snRNA-seq data collection and validation.

B) Heatmap demonstrating top canonical cell type markers for each lineage.

C) UMAP of the integrated liver atlas showing major hepatic and immune cell populations, with stacked bar plot of cell-type proportions by treatment condition and odds ratio plot for the shift in cell type proportions with hAAM therapy.

D) Violin plot and density plots illustrating canonical cell type markers in each cluster of the myeloid subset.

E) Dotplot showing gene expression in myeloid lineages

F) Stacked barplot showing nuclei proportions by treatment group and forest plot showing odds ratio plot for the shift in cell type proportions with hAAM therapy.

**Figure S2. Treatment-associated expression and enrichment in hAAM-exposed myeloid populations**

(A) Heatmaps of treatment-associated genes in *Trem2* macrophages, monocytes and Kupffer cells. Row Z-scores and fold-change annotations are shown as indicated.

(B) Enrichment among increased and decreased genes in *Trem2* macrophages, monocytes and Kupffer cells following hAAM treatment versus PBS, spanning GO Biological Process, KEGG and Reactome. Horizontal position indicates gene ratio, symbol size indicates gene hits and colour indicates $-\log_{10}$(pathway FDR).

(C) Microscope images from immunofluorescent staining of liver sections in grayscale and with merged and enlarged images showing: CLEC4F(red), TREM2 (green), and DAPI (blue), with minimal colocalisation as demonstrated (scale bars = 100µm), as well as images of staining showing CD11b (red), TREM2(green), CLEC4F (grey), DAPI (blue) with white arrows indicating CD11b/TREM2 colocalisation (below, scale bars = 10 µm).

(D) Representative images from quantification of TREM2 staining in APAP-ALI tissue in animals treated with AAM (left) vs PBS (right) with TREM2(green) and DAPI(blue) in merged images to show colocalisation.

**Figure S3. Hepatocyte-state markers and functional programmes in the hAAM study**

(A) Marker-expression heatmap across hepatocyte states with condition and state annotations.

(B) Feature plots of zonation markers *Hal*, *Cyp2e1* and *Glul*; metabolic markers *Scd1*, *Cux2*, *Angptl8*, *Thrsp* and *Fasn*; and stress-associated markers *Jun* and *Igfbp1*.

(C) Dot plot showing key marker genes for each cluster in the hepatocyte reclustering with Selected GO Biological Process terms enriched among hepatocyte-state markers.

(D) Feature plots of *Anxa2*, *Abcc4*, *Il34* and *Fosl1* and GO Biological Process enrichment among IR-Hep state markers.

**Figure S4. Characterization of EVs derived from BMDMs and AAMs from wildtype mice.**

(A) EV isolation workflow using ultrafiltration and size-exclusion chromatography.

(B) Representative BMDM and AAM morphology and expression of AAM-associated genes after IL-4/IL-13 stimulation. Morphology scale bars, 100 μm

(C) Protein and particle measurements across chromatography fractions and particle-size distributions of BMDM-EVs, AAM-EVs and medium control.

(D) Western blot analysis of CD206, GM130, HSP70, flotillin and GAPDH in AAM and BMDM lysates, EV preparations and medium controls, with representative transmission electron microscopy of EV preparations. TEM scale bar, 100 nm.

(E) Qualitative proteomic characterisation of selected EV- and macrophage-associated proteins, VENN chart showing overlap of detected proteins, and scatterplot with abundance of selected proteins in two independent EV preparations.

(F) PKH-26-labelled AAM-EV uptake by naïve BMDMs after 3 h, with control images and fluorescence quantification.

**Figure S5. In vitro functional effects of AAM-EVs in naïve BMDMs.**

(A) Dose-dependent effects of AAM-EVs on phagocytosis, shown as the fraction of pHrodo-positive cells over time (left), fluorescence intensity plotted separately by treatment (middle), and a heatmap of the fraction of pHrodo-positive cells (right). Significantly different timepoints in phagocytosis between AAM-EV treated cells vs PBS are coloured according to dose (see legend). Heatmap (right) shown for clarity demonstrates fraction of pHrodo + cells over time with each dose along the x axis.

(B) Images of PKH-26 labelled EV uptake in dose response experiment. Top panel shows cells treated with PKH-26 EVs (orange) overnight. Images shown are after 0, 5, 10 and 20 minutes after the addition of pHrodo beads (green). Cells stained with Deep Red CellMask, red; NucBlue, blue. Bottom panel shows cells treated with PBS only. *P <0.05, **P <0.01, ***P <0.001, scale bars 20 um. Twoway ANOVA with mixed effects model (n=3-6 per group).

(C) Distributions of observed $\log_2$ protein intensities and principal-component analysis of fully observed protein groups across samples. Colours indicate treatment groups.

(D) Volcano plots for AAM-EV versus PBS, BMDM-EV versus PBS and AAM-EV versus BMDM-EV. The analysis includes 8,533 protein groups; 26, 1,504 and 555 respectively meet BH FDR < 0.05. Coral and teal indicate nominal P < 0.05 with higher and lower abundance, respectively; black outlines identify BH FDR < 0.05.

The horizontal coordinate is $\log_2$ fold change and the vertical coordinate is $-\log_{10}$(nominal P).

**Figure S6. Additional dose-response and regeneration measurements after AAM-EV treatment**

(A, B) Representative p21 (A) and BrdU (B) immunohistochemistry and quantification following PBS, AAMs or 4 × $10^8$ AAM-EVs, with healthy controls.

(C) BrdU (yellow) and ERG (magenta) immunofluorescence with DAPI (cyan), and quantification of average % double-positive nuclei in animals with specified treatments. Arrows indicate double-positive cells.

(D) Serum cytokine measurements for the indicated treatments.

**Figure S7. Biodistribution and hepatic localisation of labelled AAM-EVs**

(A) Representative in vivo near-infrared imaging of healthy or APAP-injured mice given control or 4 × $10^8$ DiR-labelled AAM-EVs, at baseline and 6 and 20 h after administration, with graph showing near-infrared signal from APAP-injured mice receiving control or labelled EVs; n = 3 per group.

(B) Representative ex vivo organ imaging at 20 h with graph showing quantification of near infrared signal in harvested organs.

(C) Detection of biotin-labelled EVs in liver by DAB (left two images) and fluorescence (right two images), alongside unlabelled dye wash controls.

(D) Representative staining for DAPI (cyan), LAMP1 (yellow) and biotin-labelled EVs (magenta), with merged views and enlarged regions. Biotin detection is consistent with association with LAMP1-positive structures.

**Figure S8. Hepatic myeloid association with NHS-AF647-labelled AAM-EVs**

(A) Representative gating strategy for liver non-parenchymal cells and AF647 detection in KCs, Ly6C-high, -intermediate and -low Mo/MoMFs, and neutrophils.

(B) AF647-positive fractions from gating strategy in (A) at 1, 3 and 6 h after labelled EV administration compared with controls.

(C) Gating strategy showing separation of AF647- and AF647+ populations in live CD45+ cells. Same gating strategy as shown in (A) was then applied to each population.

(D) Graphs showing paired comparisons of AF647- and AF647+ compartments for population representation (left) and phenotype (right), with CD206 geometric mean fluorescence intensity across Ly6C subsets.

(E) Bar graphs showing population representation (left) and CD206 positivity (right) within the AF647-positive compartment at 1, 3 and 6 h.

(F) Representative Ly6C-subset (left) and CD206 (right) gates at 3 h in mice following NHS647-AAM-EV treatment (above) vs PBS (below), with bar graphs showing population proportions and CD206 geometric mean fluorescence intensity.

Individual animals and mean ± SD are shown (n = 3–5 per group, as recorded in the source legend). For two groups we used unpaired two-tailed t-tests with Holm–Šídák correction apart from D where we used paired two-tailed t-tests with the two-stage Benjamini–Krieger–Yekutieli FDR procedure at Q = 1%. For 3 groups we used two-way ANOVA. *P < 0.05, **P < 0.01, ***P < 0.001, ****P < 0.0001; ND, no discovery.

**Figure S9. GW4869 inhibition and Dicer depletion identify important mechanistic contributions of EV release and miRNA cargo**

A) Inhibition of EV secretion using GW4869 reduces particle release from day 8 AAMs in a dose-dependent manner, measured by nanoparticle tracking analysis (NTA) (top), with EV size distribution plots. Expression of canonical AAM-associated genes following treatment with increasing concentrations of GW4869 (middle).

B) Phenotype of *Dicer* flox$^{+/+}$ AAMs relative to BMDM and following incubation with Tat-Cre (left), *Dicer1* expression in *Dicer* flox$^{+/+}$ AAMs with or without Tat-Cre-mediated *Dicer1* excision (middle), and expression of selected miRNAs in AAM-EVs derived from *Dicer* flox$^{+/+}$ AAMs with or without Tat-Cre-mediated *Dicer1* excision (right).

C) Overlap of differentially represented miRNAs identified by comparison of AAM-EVs with parent AAMs, BMDM-EVs with parent BMDMs, and AAM-EVs with BMDM-EVs (left). miRNA landscape showing activation-associated enrichment (AAM-EV versus BMDM-EV; x-axis) and relative EV enrichment (AAM-EV versus parent AAM; y-axis), with point size proportional to mean abundance in AAM-EVs (CPM) (right). miRNAs detected in at least two of three AAM-EV samples with mean abundance ≥20 CPM and >2-fold enrichment in both comparisons are indicated as dual-enriched miRNAs; the four candidate miRNAs are highlighted.

D) Volcano plots showing differential miRNA abundance in AAM-EVs compared with BMDM-EVs and parent AAMs, with heatmap showing top 20 abundant miRNAs in AAM-EVs (with abundance for BMDM-EV for those miRNAs shown as comparison on the right).

E)Targeting of APAP-ALI-associated genes by the four candidate AAM-EV miRNAs (miR-149-5p, miR-181b-5p, miR-27a-3p and miR-451a), based on experimentally validated miRNA–target interactions. Genes are grouped according to functional categories associated with repair/fibrosis, DAMPs/necrosis, innate signalling, recruitment/adhesion and stress responses.

F) GO Biological Process and KEGG pathway enrichment analysis of experimentally validated targets of the four candidate miRNAs.

G) Interaction network linking the four candidate miRNAs to selected experimentally validated target genes associated with inflammatory signalling and liver injury.

**Figure S10. Characterisation and in vivo activity of GFP-AAMs and GFP-AAM-EVs**

(A) Gating and CD206 positivity of BMDMs and AAMs from C57BL/6J and MacGreen mice after IL-4/IL-13 stimulation; n = 3 independent differentiations as recorded in the source legend.

(B) EV size profiles, selected EV-associated proteins and phagocytosis measurements for MacGreen-derived EVs and the indicated controls.

(C) *GFP* transcript abundance in EVs from wild-type or MacGreen cells relative to wild-type EVs.

(D) In vivo treatment timeline: PBS, GFP-AAMs or GFP-AAM-EVs at 16 h after APAP, with analysis at 36 h.

(E) Representative H&E and Ly6G staining and quantification of necrosis and Ly6G positivity.

**Figure S11. Supporting myeloid and Kupffer-cell state analyses in the three-condition study**

(A) Myeloid marker-expression heatmap and violin plots, condition-split UMAPs, cell-type proportions with individual animals shown as dots, and treatment-effect concordance plots for MoMF1 and MoMF2.

(B) Marker-expression dotplot characterising MoMF and KC states. Dot size indicates the percentage of nuclei expressing each gene and colour indicates scaled average expression. Representative GO Biological Process terms enriched among marker genes for each MoMF and KC state. Colour indicates $-\log_{10}$(BH-adjusted P); dots are shown for terms with FDR < 0.05

(C) Concordance of treatment-associated $\log_2$ fold changes and representative KEGG and Reactome enrichment in LAM-like and core KCs.

(D) Dotplots showing cycling genes by treatment group. Dot size indicates the percentage of nuclei expressing each gene and colour indicates scaled average expression.

**Figure S12. IR-Hep treatment responses and spatial apposition of ANXA2-positive and TREM2-positive cells**

(A) Dotplot of representative IR-Hep state markers and gene-set enrichment summary of state-associated functional programmes. In the enrichment plot, colour indicates the normalised enrichment score and dot size indicates $-\log_{10}$(FDR).

(B) Spearman correlations between animal-level IR-Hep state proportions and injury-associated measurements (left), concordance of whole-compartment pseudobulk treatment effects (right). The concordance plot compares $\log_2$ fold changes for AAM versus PBS and AAM-EV versus PBS across 13,453 genes; highlighting indicates P < 0.05 in both comparisons.

(C) Representative immunofluorescence showing ANXA2-positive hepatocytes and TREM2-positive macrophages in close apposition. Individual channels and merged images are shown, with an enlarged region below. ANXA2, grey; TREM2, magenta; HNF4α, yellow; DAPI, cyan. Red arrows highlight closely apposed cells.

(D) CellChat-predicted ligand–receptor interactions from IR-Heps to TREM2 macrophages in PBS-, AAM- and AAM-EV-treated livers. Filled dots indicate condition-pooled predictions with permutation $P < 0.05$, with colour indicating the inferred interaction score. Open circles indicate $P \geq 0.05$. These P values assess predicted interactions within each condition, rather than differences between treatment groups.

#### Table S1 PCR assays and antibodies

##### A mRNA assays

| Target | Catalogue number |
|---|---|
| *PPIA* | QT00247709 |
| *B2M* | QT01149547 |
| *Chil3* | QT01549156 |
| *Arg1* | QT00134288 |
| *Mrc1* | QT00103012 |
| *Il1β* | QT01048355 |
| *IL10* | QT00106169 |
| *Inos* | QT00100275 |
| *Cd80* | QT00129787 |
| *IFNϒ* | QT01038821 |
| *Lgals3* | QT00152558 |
| *Tnf* | QT00104006 |
| *eGFP* | QT01079589 |

##### B miRNA assays

| Target | Assay identifier |
|---|---|
| cel-miR-39 | 000200 |
| mmu-miR-223-5p | 007896_mat |
| mmu-miR-124a | 001182 |
| mmu-miR-511 | 002549 |
| mmu-miR-155 | 002571 |
| hsa-miR-191 | 002299 |
| hsa-miR-146a | 000468 |

#### Table S1 Continued

##### C Antibodies and staining reagents

| Antibody or target | Supplier | Catalogue number | Dilution |
|---|---|---|---|
| CD9 | Abcam | ab307085 | 1:500 |
| HSP70 | Abcam | ab181606 | 1:500 |
| Calnexin | Abcam | ab10286 | 1:100 |
| GM130 | BD Biosciences | 610822 | 1:500 |
| GAPDH | Abcam | ab181602 | 1:1000 |
| GFP | Abcam | ab290 | 1:500 |
| CD206 | Abcam | ab64693 | 1:500 |
| BrdU | Abcam | ab6326 | 1:500 |
| HNF4α | Perseus Proteomics | PP-H1415-00 | 1:200 |
| Ly6G | BioLegend | 127602 | 1:500 |
| Ki67 | Abcam | ab16667 | 1:200 |
| ERG | Abcam | ab92513 | 1:125 |
| CLEC4F | R&D Systems | AF2784 | 1:250 |
| GPNMB | Cell Signaling Technology | 90205 | 1:50 |
| TREM2 | Merck | MABN755 | 1:200 |
| CD11b | Abcam | ab133357 | 1:5000 |
| VCAM1 | Abcam | ab134047 | 1:500 |
| ANXA2 | Cell signalling | 8235 | 1:200 |
| HNF4A | Biotechne | PP-H1415-OC | 1:200 |
| ARG1 | Cell Signalling | 93668 | 1:500 |
| BODIPY | Cambridge Biosciences | HY-W090090 | 1:500 |
| LAMP1 | Abcam | ab208943 | 1:500 |

##### D Flow cytometry antibodies

| Antibody or target | Supplier | Catalogue number | Dilution |
|---|---|---|---|
| Ly-6C PerCP-Cyanine5.5 | eBioscience | 45-5932-82 | 1:200 |
| CD11b PE-Cyanine7 | Life tech | 25-0112-81 | 1:200 |
| CD206 APC Cy7 | Biolegend | 321119 | 1:200 |
| CD45 Brilliant Violet 510 | Biolegend | 157219 | 1:200 |
| Ly-6G Brilliant Violet 421 | Biolegend | 127627 | 1:200 |
| F4-80 PE | Biolegend | 111603 | 1:200 |
| MHC II Brilliant Violet 741 | Biolegend | 107643 | 1:200 |
| CD64 BV605 | Biolegend | 139323 | 1:200 |
| GFPGFP | Abcam | ab13970 | 1:200 |
| CD206-PE | Biolegend | 141705 | 1:200 |
